# Complex genetic architecture underlying the plasticity of maize agronomic traits

**DOI:** 10.1101/2022.01.18.476828

**Authors:** Minliang Jin, Haijun Liu, Xiangguo Liu, Tingting Guo, Jia Guo, Yuejia Yin, Yan Ji, Zhenxian Li, Jinhong Zhang, Xiaqing Wang, Feng Qiao, Yingjie Xiao, Yanjun Zan, Jianbing Yan

**Affiliations:** National Key Laboratory of Crop Genetic Improvement, Huazhong Agricultural University, Wuhan, 430070, China; Gregor Mendel Institute, Austrian Academy of Sciences, Vienna BioCenter, 1030, Vienna, Austria; Institute of Agricultural Biotechnology, Jilin Academy of Agricultural Sciences, Changchun, 130033, China; Hubei Hongshan Laboratory, Wuhan, 430070, China; College of Life Sciences, Sichuan University, Chengdu, 610065, China; Institute of Agricultural Sciences of Xishuangbanna Prefecture of Yunnan Province, Jinghong, 666100, China; Umeå Plant Science Center, Department of Forestry Genetics and Plant Physiology, Swedish University of Agricultural Sciences, Umeå, 90736, Sweden

**Keywords:** Complex traits, Phenotype plasticity, QTL by environment interaction, Crop improvement, *Zea mays*

## Abstract

Phenotypic plasticity is the property of a given genotype to produce multiple phenotypes in response to changing environmental conditions. Understanding the genetic basis of phenotypic plasticity and establishing a predictive model is highly relevant for future agriculture under changing climate. Here, we report findings on the genetic basis of phenotypic plasticity for 23 complex traits using a maize diverse population, planted at five sites with distinct environmental conditions and genotyped with ~ 6.60 million SNPs. We found that altitude-related environmental factors were main drivers for across site variation in flowering time traits but not plant architecture and yield traits. For 23 traits, we detected 109 QTLs, of which 29 was for mean, 66 was for plasticity, and 14 for both parameters, besides, 80% of the QTLs were interreacted with the environment. The effects of several QTLs changed in magnitude or sign, driving variation in phenotype plasticity, and we further experimentally validated one plastic gene *ZmTPS14.1* whose effect was likely mediated by the compensation effect of *ZmSPL6* which was from the downstream pathway probably. By integrating genetic diversity, environmental variation, and their interaction in a joint model, we could provide site-specific predictions with increased accuracy by as much as 15.5%, 3.8%, and 4.4% for DTT, PH, and EW, respectively. Overall, we revealed a complex genetic architecture involving multiallelic, pleiotropy, and genotype by environment interaction underlying maize complex trait mean and plasticity variation. Our study thus provided novel insights into the dynamic genetic architectures of agronomic traits in response to changing environments, paving a practical route to precision agriculture.

## Introduction

Upon climate change, plants display plastic response, where a single genotype produces multiple phenotypes through changes in gene expression, physiological and morphological levelsl^1,2^. Such plastic response (phenotype plasticity) was also described as genotype by environment interaction (G-by-E)^3–5^, with organisms changing their performance across environments, releasing heritable variation^6–9^ that are highly relevant in complex trait variation and adaptation^4,10–12^. In the context of crop breeding, one strategy is to minimize plasticity or G-by-E interaction by using the best linear unbiased prediction value (BLUP), making developed cultivar broadly applicable to a wide range of environments^13^. Alternatively, performance could be maximized in individual environments by enriching site-specific beneficial alleles that are either neutral or unfavourable at other sites^12,14^. This is similar to what natural selection have acted on wild populations, where local adaptation has resulted in genotypes with optimized phenotypes at their native environments that are often maladapted in new environments^15–18^.

Increased plasticity may represent the future of crop breeding and biodiversity management in the light of climate change, as such strategy confers high resilience genotypes for future challenges while achieving optimal phenotype locally. To achieve this goal, efforts have been made to study the genetic architecture of plasticity ^19–22^ and dissect the underlying QTLs ^4,11,23–26^. Studies in maize have revealed both similarity and difference in the genetic architectures of trait mean and plasticity^24,25^, suggesting breeders could manipulate trait mean and plasticity semi-independently to meet the challenge of feeding the growing population. Further investigations demonstrated the role of plastic QTLs in heterosis and adaptation from tropical to temperate zone, paving the way to genomic-promised crop improvement by manipulating the phenotypic plasticity^27,28^.

Despite the insights gained through these efforts, several questions remain elusive. First, there is a lack of understanding of the dynamics of complex traits genetic architectures across environments, such as the impact of specific environmental factors on range-wide complex trait variation, how dynamic are the genetic architectures of agronomic traits over major production zone? What alleles are favoured at each production site? Whether they have genetic effects on multiple traits with antagonistic pleiotropy? How much genetic gain could be achieved by exploiting these alleles?

Second, in Fisher’s decomposition of phenotype mean, the environmental effect is a combinatory effect from multiple environmental factors, such as temperature, day length, and soil conditions, etc. With an increased ability to quantify air and soil conditions using developments in remote sensing, it is of great interest to decompose the combinatory environment effects into effects from concrete environmental factors and study their impact on complex trait variation and prediction. Last but not least, plasticity was often treated as a composite index^19–22^, neglecting the fact that plasticity is environment-dependent, being variable when quantified using different combinations of environments. With a growing number of environments that we could investigate, it is worthwhile to differentiate plasticity quantified using an overall index and refine plasticity measures from specific combinations of environments.

To provide a deeper insight into these questions, we developed the Complete-diallel plus Unbalanced Breeding-derived Inter-Cross (CUBIC) population of 1404 advanced inter-cross lines from 24 representative breeding founders^29^ and studied the variation of 23 key agronomic traits at five sites spanning China’s major summer maize production zone (Fig. 1A) from northeast at Jilin (JL; N 43° 42’, E 125°18’) to central plains at Henan (HN; N 35° 27’, E 114° 01’). We revealed major contributions from the latitude-related environmental factors to across site phenotypic variation for flowering time traits but not for others. And we dissected the within and across environment variation to 109 QTLs with complex genetic architectures involving multiallelic, pleiotropy, and genotype by environment interaction. In particular, we found that extensive QTL by environment interaction and dynamic in mean QTL effects across environments was driving the variation in phenotype plasticity. A joint model with site-specific predictions and higher accuracy was developed by integrating genetic diversity, environmental variation, and their interaction, paving a way to genomics-directed maize improvement.

**Fig. 1.**
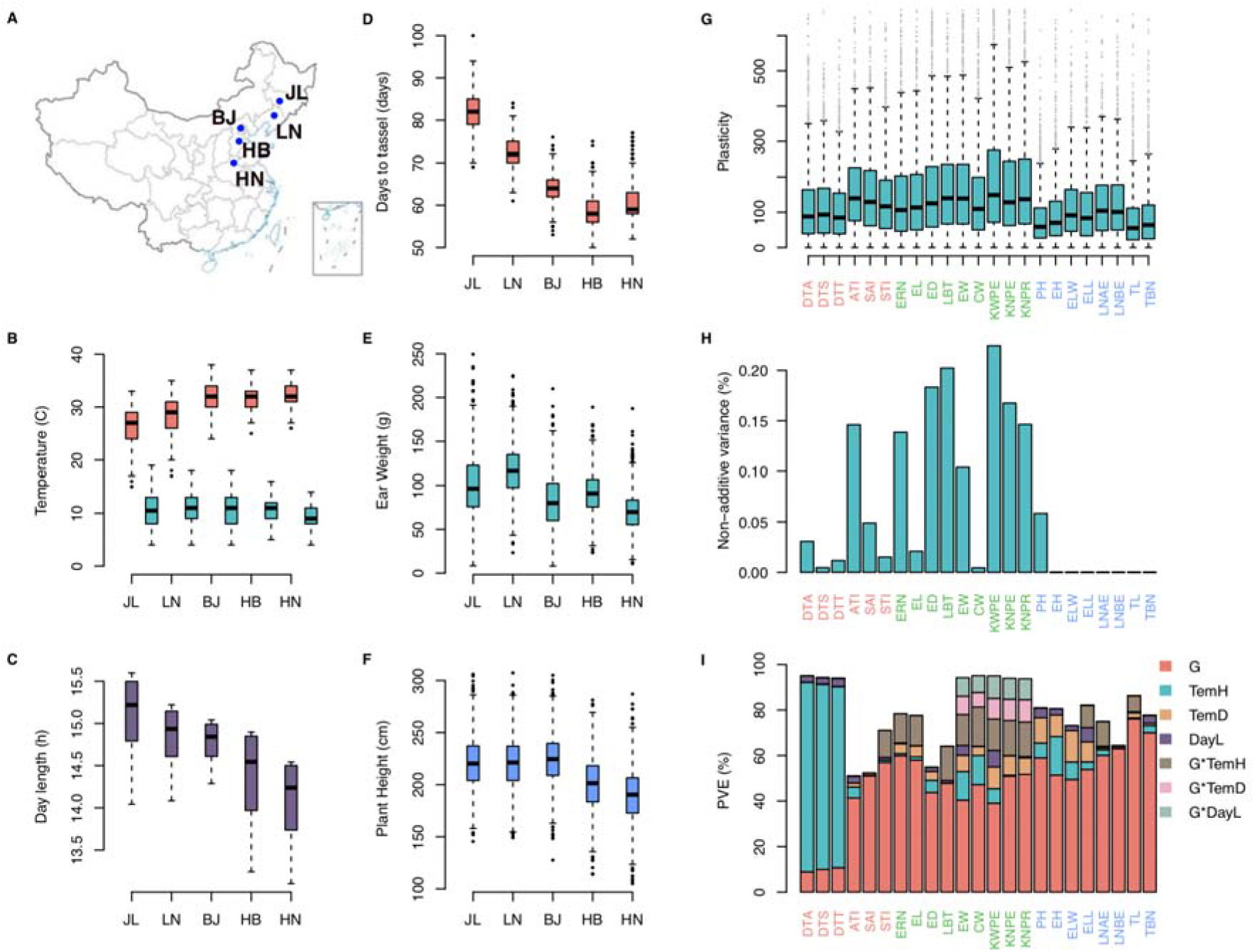
Environmental variation across China’s major summer maize production zone and their impact on the across-site variation of maize complex traits. **A)** The five surveyed sites spanning China’s major maize production zone, where 23 agronomic traits were phenotyped for 1404 inbred lines. **B)** Boxplot illustrating the highest daily temperature (TemH; coloured in cyan) and daily temperature difference (TemD; coloured in tomato) from sowing to flowering at the five sites. **C)** Boxplot of the day length (DayL) from sowing to flowering at the five sites. **D)** Boxplot of Days to tassel (DTT; coloured in tomato), **E)** Ear weight (EW; coloured in green) and **F)** Plant height (PH; coloured in cyan) measured at the five sites. **G)** Boxplot of the phenotype plasticity measured as a coefficient variation of the rank across s (Materials and Methods). The 23 traits (labelled in x-axis) were grouped into 3 categories, flowering traits highlighted in tomato, plant architecture traits labelled in green and yield traits labelled in blue. **H)** Bar plot of the proportion of non-additive variance (differences between broad-sense heritability, capturing the additive and non-additive effect, and narrow-sense heritability, capturing only the additive effect). Each vertical bar represents a trait, and the height of the bar is proportional to the difference between corresponding broad and narrow-sense heritability. **I)** Contribution from genotype, the three environmental factors (TemH, TemD and DayL), and their interactions to the across-site variation of the 23 agronomic traits. Each vertical bar represents a trait with the corresponding trait name labelled in x-axis. The coloured segments within each bar represent the contribution from G, TemD, TemH, DayL, and their interaction with G as indicated in the legend. The height of the segment is proportional to the variance explained (PVE) by the corresponding variance component.

## Results

### The impact of clinal variation in environmental factors on the mean and plasticity of 23 maize complex traits

We surveyed the performance of 23 traits across five sites spanning Chinese major summer maize production zone with longitudinal variation from E114° 01’ (Henan; HN) to E125° 18’ (Jilin; JL) and latitudinal variation from N 43° 42’ (JL) to N 35° 27’ (HN; Fig. 1A). Across the five sites, daily highest temperature (TemH), daily temperature difference (TemD), and day length (DayL) varied significantly (Fig. 1B, C). Nearly all the traits (22 out of 23, except for Leaf number below ear, LNBE) were significantly correlated with latitude at the five sites, suggesting a general contribution from spatially variable environmental factors to maize agronomic trait variation (Fig. 1D-F; Figure S1; Table S1, S2). Flowering time traits (days to tassel DTT; days to silking, DTS; days to anthesis, DTA) displayed the strongest latitudinal variation with trait median measured at JL being ~1.5 times larger than that at HN (Figure S1; Table S2). Unlike flowering time, clinal variation in plant architecture traits (Plant height, PH; Ear height, EH; Ear leaf width, ELW) and yield traits were weaker, being more distinctive between the northern (JL, LN, and BJ) and southern (HB and HN) sites (Fig. 1B, C; Figure S1).

To explore how the 23 traits responded to the across-site environmental perturbation, we first rank-transformed each trait measured at individual sites and quantified the phenotype plasticity as coefficient variation of rank (VarR) across the five sites. All the 23 traits displayed variation in phenotype plasticity (Fig. 1G), and yield traits were more plastic than flowering time and plant architecture traits. Contributions from environment (E) and genotype by environment interaction (G-by-E) varied significantly among the three categories of traits. For example, TemH was the major driver for (median = 84.2%) across-site variation of flowering time traits (DTT, DTA, and DTS; Fig. 1I), while its contribution to the variation of remaining traits was much lower (Fig. 1I; media=9%). In contrast, G-by-E made a higher contribution (median = 32.8%; Fig. 1I) to the across site variation of yield traits, being consistent with the observation that the proportions of non-additive variance for yield traits were also higher than that for flowering and architecture traits (Fig. 1H). Altogether, these results illustrated a general contribution from environment factors (TmpD, TmpH, and DayL) and their interaction with genotype to the variation of maize complex traits, where the contribution from G-by-E was more prominent for yield traits, indicating the importance and potential value of studying plasticity for yield improvement.

### Dynamic and complex genetic architecture underlying maize agronomic traits mean and plasticity

For each of the 23 traits, we derived two types of measures to quantify the phenotype plasticity, where type I included 10 measures^30,31^ calculated as pairwise difference among five sites to capture specific plasticity (SP), and type II included 4 measures representing overall plasticity (OP): coefficient of variation from raw (CV) ^30^, rank transformed data (VarR) ^30^, second principal component (PC2) ^30^, and Finlay–Wilkinson regression (FWR) ^32^ (Figure S2; Materials and Methods). Together with trait mean value from five sites (Mean) and BLUP, these four types of measures (SP, OP, Mean, and BLUP) were used to scan for QTLs underlying trait mean and plasticity, using genome-wide association analysis with 6.6 M genetic polymorphisms (Materials and Methods). In the following section, we first illustrated the results from DTT as an example and then expanded to results from all 23 traits. Hereafter, the 4 types of measures were referred to as DTT_BLUP_, DTT_x_ (mean measured at site X), SP-DTT_x-y_ (Specific plasticity measured as DTT_x_-DTT_y_, X and Y was site name), and OP-DTT_z_ (Overall plasticity calculated using method z, z was described in Materials and Methods).

#### Loci associated with the variation of mean and plasticity measures for days to tassel – Dynamic QTL effects across environments lead to variation in plasticity

A total of 15 QTLs were identified, including 11 QTLs for SP/OP-DTT and 7 QTLs for DTT_mean_/DTT_BLUP_ with 3 overlaps (Fig. 2A-D, QTLs were obtained by grouping independent SNPs within defined physical distance, Materials and Methods; Table S3, S4, S5). A majority of the QTLs were detected for DTT_mean_ and SP/OP-DTT, while only 2 QTLs were detected for DTT_BLUP_, highlighting the added value to analyse DTT_mean_ and the derived plasticity measurements individually (Fig. 2B). By contrasting genetic effects of QTLs across sites, two types of QTLs, whose effects changed in magnitude or direction, were detected with significant contribution to the variation of DTT plasticity. For example, different genotypes of QTL8 (chromosome 5: 6,462,711 bp, Fig. 2A-C, E, 667.2 kb upstream of *ZmPHYC2*, GRMZM2G129889, a homology of *Arabidopsis thaliana PHYC*^33^) showed a significant phenotypic difference for DTT_HN_ (P = 1.2 x 10^-7^; Fig. 2D) and the specific plasticity measures, calculated as the difference between HN to the other sites (e.g. DTT_HN-BJ_; P = 5.3 x 10^-8^; Fig. 2E), but had no effect at the remaining DTT mean and plasticity measurements (Fig. 2C-E), indicating changes in the magnitude of genetic effects contributed to the variation of DTT plasticity. In contrast, QTL14 (chromosome 9: 35,126,793 bp, Fig. 2A-C, F, 508.5 kb upstream of *CONZ1*, GRMZM2G405368, a homology of Arabidopsis thaliana *CO*^34^), was exclusively detected for several DTT plasticity measurements but not for any of the DTT_mean_ and DTT_BLUP_. The genetic effects of QTL14, however, changed direction from positive (DTT_HN_, Additive effect = 0.6 ± 0.2 days; P =1.7 x 10^-3^; Fig. 2F) to negative (DTT_HN_, Additive effect = −0.5 ± 0.2 days; P = 7.3 x 10^-3^; Fig. 2F), leading to significant association with specific plasticity, DTT_HN-BJ_ (Additive effect = 1.1 ± 0.2 days; P = 6.3 x 10^-12^; Fig. 2D) and overall plasticity, DTT_pc2_ (P = 4.1×10^-9^), The detection of such loci highlighted the increased power by analysing plasticity measurements.

**Fig. 2.**
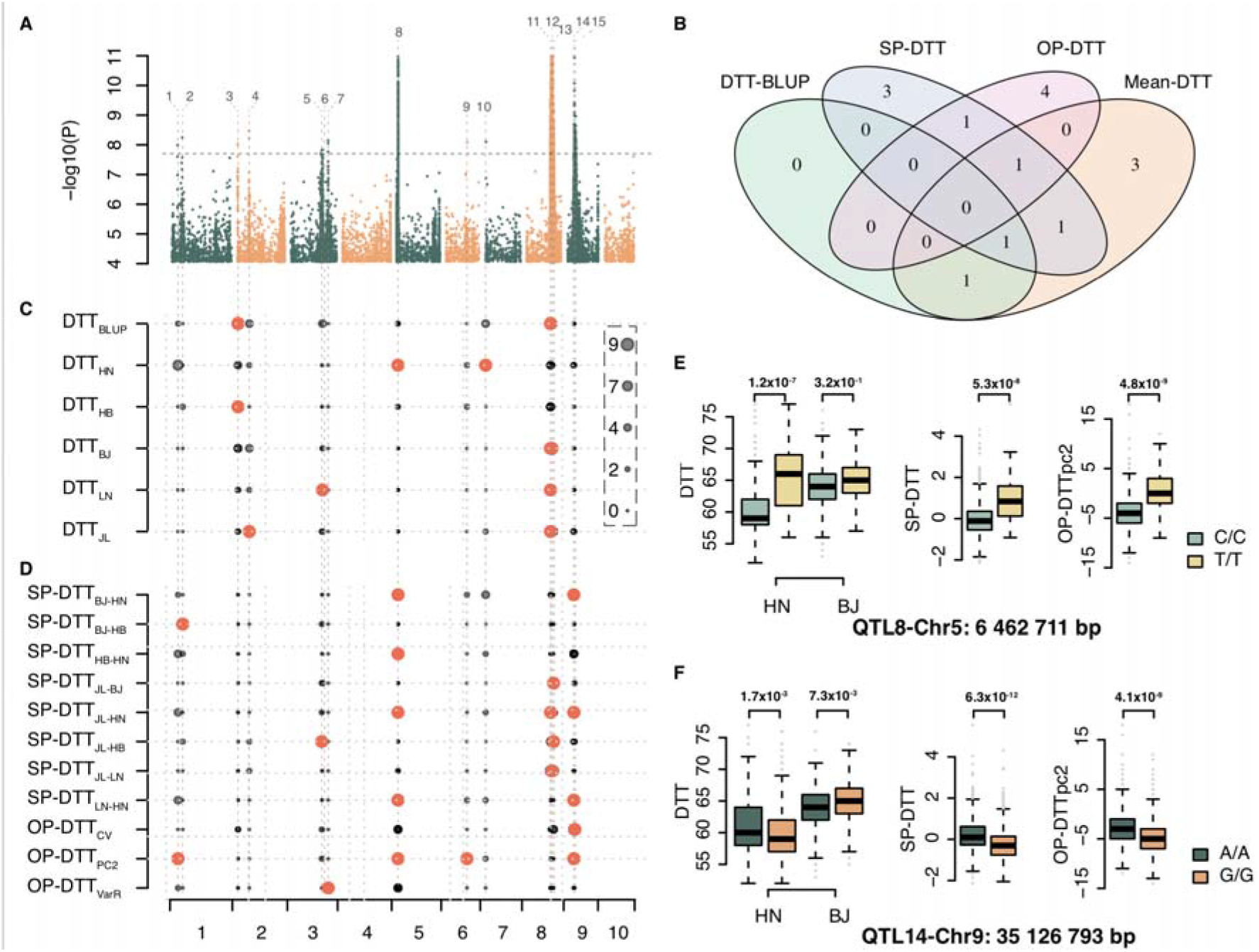
Summary of the QTLs associated with the mean and plasticity measures for days to tassel (DTT). **A)** Manhattan plots overlaying genome-wide association scan results for the 20 mean and plasticity measurements for DTT. The black horizontal dashed line indicated the Bonferroni-corrected genome-wide significance threshold derived as 0.05/Me (Me is the effective number of independent SNPs; Materials and Methods), and the vertical dashed black lines indicate the position of detected QTLs, labelled from 1 to 15. **B)** Venn diagram illustrating the overlap of QTLs detected for the 4 types of DTT measurements. **C)** QTLs associated with the DTT means measured at five sites and the DTT_BLUP_ (y-axis). Each dot represents a SNP and the size of the dot is proportional to its −Log10 p value as indicated in the legend on the right. Loci with p-value passed genome wide significance threshold were coloured in tomato. **D)** QTLs associated with the DTT plasticity measurements (labelled in the y-axis). **E) and F)** Genotype-to-phenotype maps, highlighting the increased power to detect additional loci by analysing plasticity measurements, for DTT_HN_, DTT_BJ_, DTT_HN-BJ_, and DTT_pc2_ at two QTLs, one at chromosome 5: 6,462,711 bp and a second one at chromosome 9: 35,126,793 bp.

Altogether, these results indicated that changes in magnitude and/or signs of genetic effects across sites caused variation in plasticity, which could be detected by GWAS on SP and OP measurements. The changing genetic effects highlighted the role of QTL by environment interaction in the variation of complex trait mean and plasticity.

#### Loci associated with the variation of remaining traits– A complex genetic architecture involving multiallelic, pleiotropy, and genotype by environment interaction underlay maize complex trait variation

For the 23 traits, we identified 109 QTLs for the 4 types of measurements (Fig. 3A; Figure S3; Table S3, S4, S6), which overlapped partially, with 1.8%, 34.9%, 19.3%, and 21.1% of the QTLs being unique to BLUP, SP, OP and Mean measurements, respectively (Fig. 3B). As has been illustrated in the previous section, QTLs associated with SP measurements likely changed their genetic effects in sign or direction (Fig. 2E, F). This was supported by testing the interaction between the detected QTLs and the five sites, where 80.0% of the QTLs were found to be significantly interacting with the sites (Table S7; Materials and methods). This demonstrated the dynamic genetic effects of mean QTLs across sites and highlighted partial overlap for QTLs regulating mean and plasticity as reported in the previous studies ^24,25,27^.

**Fig.3.**
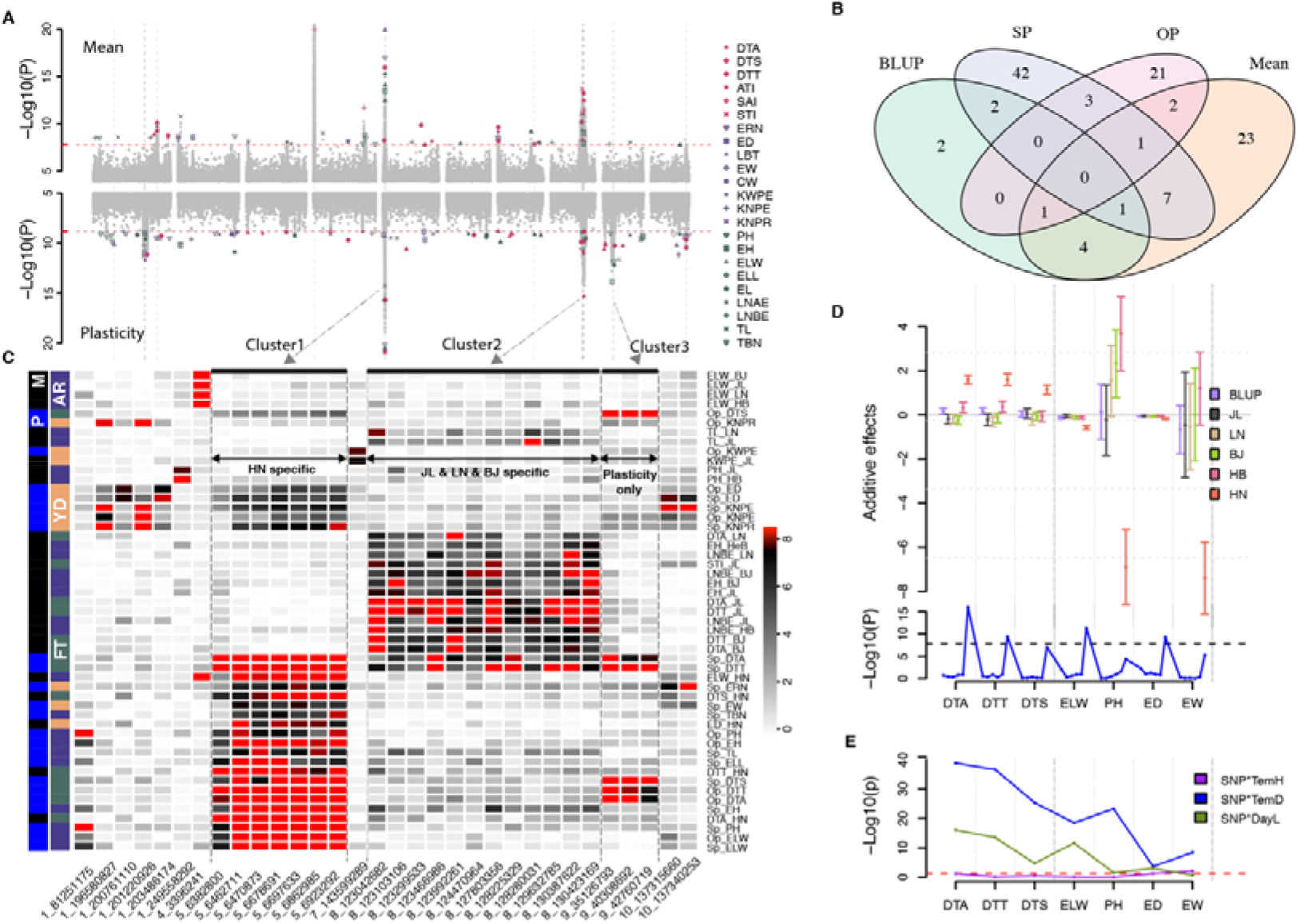
Association results of both mean and plasticity for all the 23 traits. **A)** Manhattan plots of GWAS from all scans, with upper panel for means and lower panel for plasticity measurements. The red horizontal dashed lines indicate the Bonferroni-corrected genome-wide significance threshold. The vertical dashed grey lines highlight the site of 32 SNPs associated with more than 2 measurements. **B)** Venn diagram illustrating the overlap of QTLs detected for the 4 types of measurements. **C)** A heatmap illustrating the p values of the 32 SNPs detected for more than 2 measurements (Here, SNPs were used instead of QTLs, as one QTL sometimes includes multiple statistically independent SNPs that are physically close to each other). Each cell represents the −Log10 (p values) of a particular SNP (x-axis) associated with a specific trait (y-axis on the right). The outer index on the left side marks the mean (M, in black) or plasticity (P, in blue) of the traits. The inner index marks the corresponding trait types: plant architecture (AR; in purple), flowering time (FT; in olive-green), and yield (YD; in orange). For each trait, only the lowest p-values were indicated for either specific plasticity (SP) or overall plasticity (OP), labelled as SP-trait or OP-trait. **D)** The Additive effects varied across sites exampled for cluster 1 (chromosome 5:6 462 711 bp) on multiple traits. The traits were separated by the dashed vertical lines and labelled in x-axis, and for each trait the measurements for BLUP, individual sites were ordered (from left to right) as indicated in the colour legend (from top to bottom). Median and standard error were shown with the middle point and error bars. The corresponding GWAS p values were illustrated in the lower panel. **E)** The p-values testing the interaction between this SNP (chromosome 5:6 462 711 bp) and the 3 environmental factors.

One QTL, spanning 540 kb from chromosome 5: 6,382,800 bp to 6,923,292 bp, involved 7 statistically independent SNPs (Cluster 1 in Fig. 3A, C, QTL 8 in Fig. 2A) and was detected for multiple trait means and plasticity measures at HN. A detailed exploration showed that multiple haplotypes were underlying this region with each of the 7 SNPs tagging unique haplotype (Figure S4), suggesting that the 24 founders carried different functional variants. Moreover, each of the 7 SNPs was simultaneously associated with multiple trait means at HN, including flowering time (DTT_HN_ and DTA_HN_), plant architecture trait (the ear leaf width, ELW_HN_), and multiple SP measurements (SP-DTT, SP-DTA, SP-DTS, SP-PH, SP-EW) at genome wide significance (Fig. 3C, D). Moderate association to the mean and plasticity measurements for yield and plant architecture traits were also found at a relaxed significance threshold (P = 6.0 x 10^-6^ for EW_HN_ and P = 5.1 x 10^-5^ for PH_HN_; Fig. 3D), indicating this region was highly pleiotropic. Notably, the genetic effect of this QTL was unique to HN for all the associated traits, where the “TT” genotype increased DTT, DTA, DTS and “CC” decreased ED, EW, ELW, and PH at HN but not at other sites, likely due to interaction with temperature (Especially TemD) rather than DayL (Fig. 3E; Figure S5), providing an ideal candidate for targeted breeding application at HN.

A second cluster, spanning 7.4 Mb from chromosome 8: 123,042,682 bp to 130,423,169 bp, showed both allelic heterogeneity and pleiotropic effect on multiple flowering and plant architecture traits (Cluster 2 in Fig. 3A, C; Figure S6). However, their genetic effects were unique to the three northern sites (JL, LN, and BJ; Fig. 3C), except for LNBE_HB_. Compared with cluster 1, whose effects were unique to HN, such regional effects on multiple northern sites may have led to the detection of this QTL for multiple sites BLUP measurements.

A third cluster (Cluster 3 in Fig. 3A, C) was found contributing exclusively to the variation of plasticity measurements for all the flowering time traits (DTT, DTA, and DTS) due to the change of additive effects from negative to positive (Fig. 2F).

Altogether, these results illustrated a complex genetic architecture involving multiallelic, pleiotropy, and genotype by environment interaction underlying maize complex trait variation. The detection of QTL unique to HN and the three northern sites demonstrated a variable genetic architecture of maize complex traits across sites possibly due to clinal variation in QTL effects.

### The possible molecular basis of phenotype plasticity

Previously, we linked *ZmTPS14.1* (GRMZM2G068943, chromosome 8: 123,129,008 bp to 123,140,283 bp) to variation of flowering time mean^35^, which was located inside the QTL on chromosome 8 (cluster 2 in Fig. 3, chromosome 8: 123,042,682 bp to 130,423,169 bp). Here, this QTL was simultaneously associated with mean and plasticity variation of multiple traits, including DTT, DTA, DTS, ATI, STI, SAI, LNAE, and LNBE at genome-wide (P =1.53 x 10^-8^) or suggestive significance threshold (P =1.00 x10^-5^) and the tagging SNPs were interacting with all 3 environmental factors, suggesting a general contribution from QTL by environmental factor interaction to variation in phenotype plasticity.

To experimentally validate and evaluate the plasticity effects of *ZmTPS14.1*, we planted the knock-out lines of *ZmTPS14.1* obtained in a previous sudty in Jilin (JL, North China, N 43° 30’, E 124° 49’) and Hainan (HaiN, South China, N 18° 34’, E 108° 43’) and compared the measured flowering time phenotypes. In consistent with the association results, the female flowering time (DTS) of knock-out lines was earlier in HaiN but not significantly changed in JL compared to wildtype lines (Fig. 4A; Figure S7A; Table S8). To further explore the underlying molecular basis, we analysed an in-house time-course transcriptome dataset generated from reference accession B73 under long-day and short-day conditions (Fig. 4B). The expression of *ZmTPS14.1* under both day-length conditions changed in the same direction along the time course (Fig. 4B), suggesting there was no day-length dependent expression response for *ZmTPS14.1*.

**Fig. 4.**
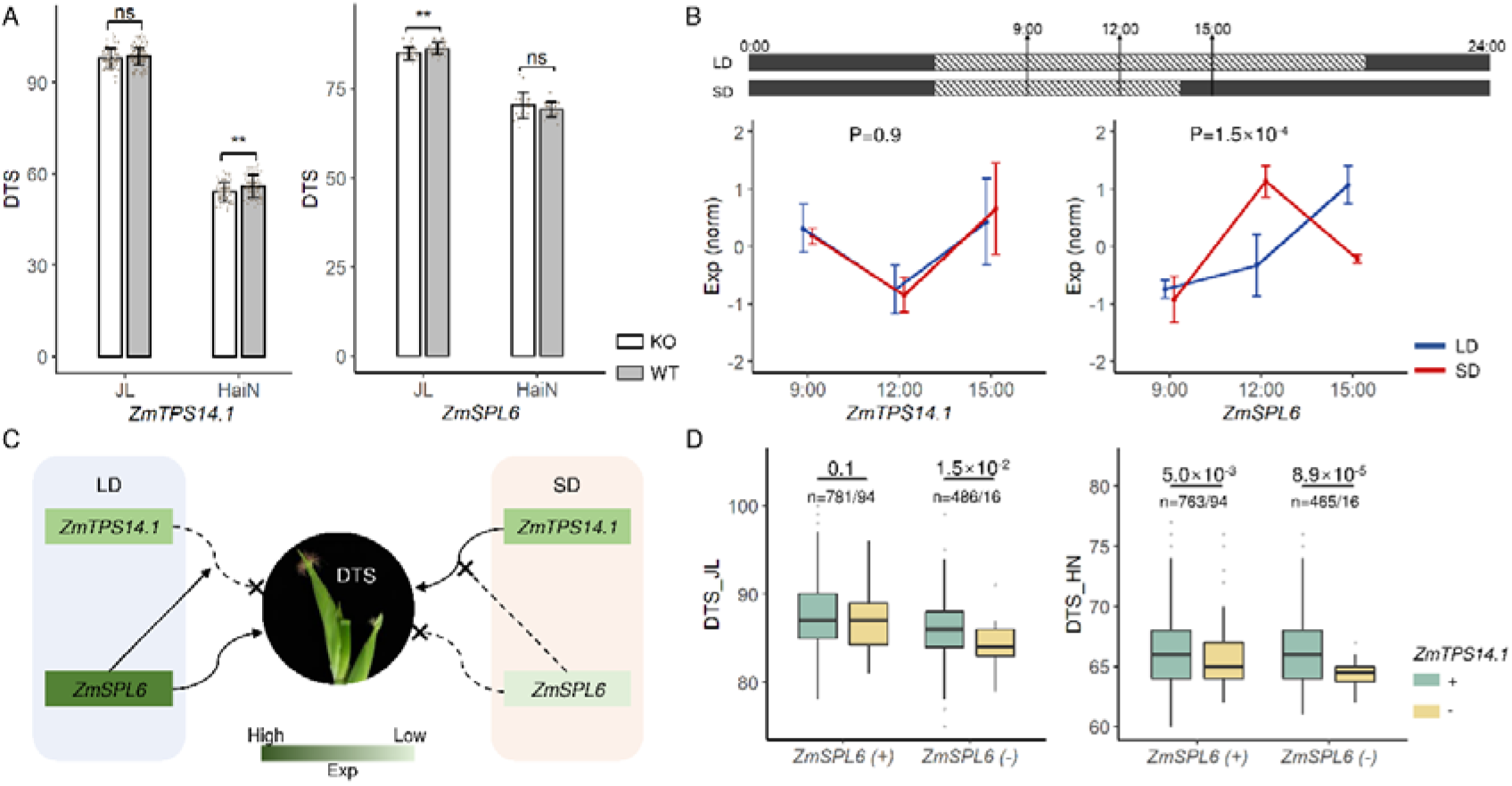
The interaction between *ZmTPS14.1* and *ZmSPL6* reveals the genetic basis of phenotype plasticity of flowering time. **A**) Phenotype (DTS; days to silking) of knock-out lines and wild type of *ZmTPS14.1* and *ZmSPL6* at two field plantations, one plantation at JL represents Jilin (N 43° 30’, E 124° 49’), and another one at HaiN represents Hainan (N 18° 34’, E 108° 43’). Error bars represent standard deviation. ** indicate P values < 0.01 by Student’s t-test. “ns” means no significance. **B)** The sampling diagram of the time-course experiment in B73 under long-day (LD) and short-day (SD) conditions. The black area represents dark time and the dotted-line area represents light time. Leaf tissues were harvested at three time points (9:00, 3 hours of light; 12:00, 6 hours of light; 15:00, 9 hours of light/ 1 hour of dark). The expression pattern of *ZmTPS14.1* and *ZmSPL6* at three time points under the long-day condition (LD, blue) and the short-day condition (SD, red) were shown. The y-axis represents gene expression, which was obtained from standardization of raw reads counts then z-score normalization. Error bars represent standard error. **C)** The proposed compensation interaction model between *ZmSPL6* and *ZmTPS14.1. ZmSPL6* expressed highly in the long-day condition which could promote female flowering but its expression suppressed in the short-day condition (SD) showed no effect for flowering. And the knockout lines of *ZmTPS14.1* showed the flowering time difference in the short-day condition, but not in the long-day condition because of the compensation effect of *ZmSPL6.* **D)** The phenotype (DTS in JL and HN) comparison between two alleles of *ZmTPS14.1* (chromosome 8: 123,138,468 bp; C/C genotype +; T/T genotype → -) in the different allele background of *ZmSPL6* in LD (JL) and SD (HN) conditions (chromosome 3: 159,420,596 bp; G/G genotype → +; T/T genotype → −). P-values were obtained by Student’s t-test.

As has been proposed that plastic response may involve developmental switch genes^36^, we explored whether plastic effects of genes at the center of the regulatory pathway were mediated or interacted with downstream genes. Therefore, we evaluated the expression of candidates downstream of *ZmTPS14.1. ZmTPS14.1* encodes Trehalose-6-phosphate synthase (TPS), which converts glucose-6-phosphate into Trehalose-6-phosphate (T6P), regulating vegetative development and flowering by miR156/SPL pathway^37^. The expression of *ZmSPL6* (GRMZM5G878561), an SPL family member downstream of *ZmTPS14.1*, showed a significant expression pattern difference in response to long/short day length (Fig. 4B). Meanwhile, the knock-out lines of *ZmSPL6* showed earlier female flowering in JL but no significant change in HaiN compared to wildtype lines (Fig. 4A; Figure S7B; Table S8), suggesting day length was an important factor for the plastic effect of *ZmSPL6.* Thus, we proposed a compensation mechanism from *ZmSPL6* to *ZmTPS14.1* in DTS plasticity (Fig. 4C). In the long-day condition, the continuous expression increase of *ZmSPL6* could make up for the knockout effect of *ZmTPS14.1*, resulting in no phenotypic difference between the knock-out lines of *ZmTPS14.1* and wildtype (Fig. 4A). But no such compensation appeared in the short-day condition, thus we observed the phenotype difference between knockout and wildtype lines of *ZmTPS14.1* in the short-day condition (Fig. 4A, C). This compensation mechanism was also reflected in the CUBIC population (Fig. 4D). In the long-day condition (JL), *ZmTPS14.1* (chromosome 8: 123,138,468 bp) showed significant association (P = 1.5 x 10^-2^) with DTS in the TT allele background of *ZmSPL6* (-), but not significant in GG allele background of *ZmSPL6* (+). And in the short-day condition (HN), the significant association between *ZmTPS14.1* and DTS was detected in both *ZmSPL6* (-) (P = 8.9 x 10^-5^) or *ZmSPL6* (+) (P = 5.0 x 10^-3^) backgrounds (Fig. 4D).

### Accounting for dynamics in genetic architecture improved complex traits prediction across environments

We evaluated the potential of integrating genetic diversity, environmental variation, and their interaction in complex trait prediction by jointly modelling genotype, environment, and their interaction (referred to as GEAI model, Materials and Methods). Two cross-validation schemes were considered. In the first case, we explored the predictability on untested lines at any of the five sites by using all the lines phenotyped at the five sites using 5-fold cross-validation. Compared with the GBLUP, with a universal prediction for all sites, our model not only provided site-specific predictions but also increased prediction accuracy for a majority of traits and sites (83.0% of all traits and sites; Table S9, Materials and methods). The averaged prediction accuracy for DTT, PH and EW increased by 5.3%, 1.2%, and 1.8%, respectively, and the increase in prediction accuracy was more pronounced at HN (increased by 15.5%, 3.8%, and 4.4% for DTT, PH, and EW, respectively, Fig. 5A-C).

**Fig. 5.**
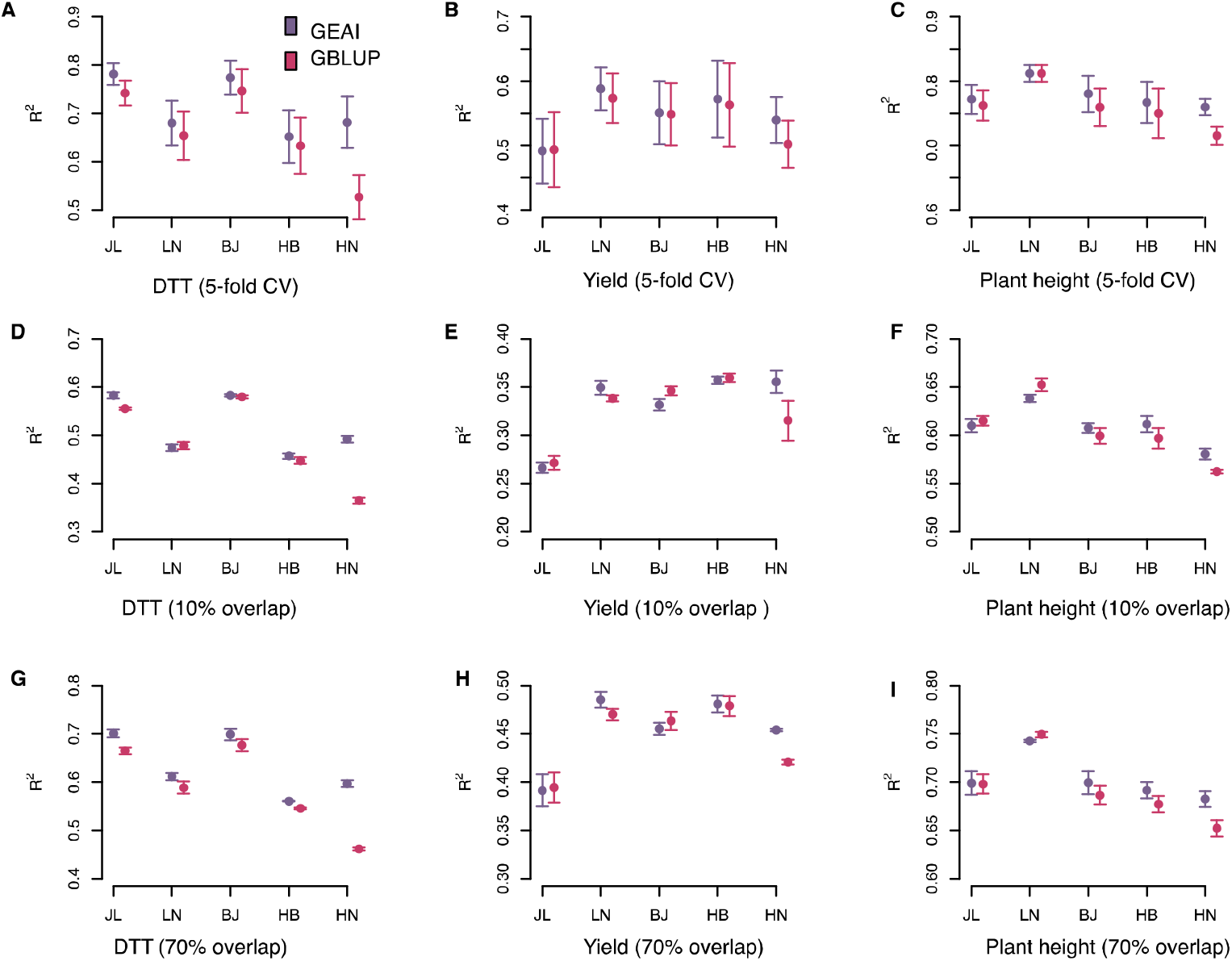
Performance of GEAI model in site-specific complex trait prediction. **A)** Predictability of untested lines at any of the five sites by using all the lines phenotyped across the five sites as training data for **A)** DTT, **B)** EW, and **C)** PH. Prediction accuracy for untested lines at any of the five sites for **D)** DTT, **E)** EW, and **F)** PH when 10% of the lines were phenotyped at all five sites and remaining lines were only phenotyped at one of the five sites. Prediction accuracy for **G)** DTT, **H)** EW, and **I)** PH when 70% of the lines were phenotyped at all five sites and the remaining lines were only phenotyped at one of the five sites.

In the second case, we explored a serial of more challenging designs, in which only a core set of lines (10%- 70%) were phenotyped across five sites and the interest was to predict the performance of unphenotyped lines at each site. It was very encouraging to see that our GEAI model showed higher accuracy for almost all the traits and sites. For example, at 10% overlap, our GEAI model outperformed GBLUP predictions (P = 4.0 x10^-3^) by 3.2% on average and increased the prediction accuracy at four out of five sites by 1.0%-12.7% for DTT, and the averaged accuracy was increased to 4.6%. At 70% overlap, the increase in accuracy at each site was larger than at 10% overlap (1.5%-13.5%; Fig. 5D-I). As the number of lines phenotyped at all sites increased from 10% to 70%, both the averaged accuracies and site-specific accuracies increased (Figure S8). Overall, our study highlighted the potential of intergrading QTL by environment interaction in understanding complex traits variation and predictions.

## Discussion

Here, by surveying the performance of a genetically diversified population across China’s summer maize major production zone, we were able to quantify contributions from specific environmental factors to the variation of 23 complex traits, detect plastic QTLs, and provide site-specific complex traits prediction model with higher accuracy.

### Contribution from environmental factors to maize complex traits mean variation and plasticity variation

Plants time their vegetative and reproductive growth in response to changes in seasonal cues, such as winter temperatures (vernalization) and day length (photoperiod)^38^. Although many studies have emphasized the importance of photoperiod to flowering time regulation, the temperature is a key determinant of flowering time^39–41^, and significantly stronger correlations between seasonal transcriptome and temperature than those with day length^42^ were reported. In consistent with these reports, we found that TemH had a considerable high contribution to the across-environment variation of flowering traits, while very little contribution from TemD, DayL, G-by-TemH, or G-by-DayL were found. In contrast, yield traits were influenced by a combination of TemH, TemD, DayL, and their interactions with genotype. A possible explanation is that both photosynthesis and respiration losses, mainly determining the crop yields, are sensitive to temperature and day length^43^, and previous studies have shown that temperature and day length could also affect days to maturity, rate of yield accumulation, and harvest index^44^.

Unfortunately, soil conditions, such as pH, soil temperature, water, and nutation content were not available in our study, limiting our ability to provide broader insight into the impact of specific environmental factors on complex trait variation. The genotype to filed (G2F) Maize project^45^, one of the ongoing efforts aiming at compensatively surveying the environmental factors and performance of diversified population across a large number of field plantations, would be of great importance to characterise the role of specific soil factors on complex traits variation.

Here, we quantified phenotype plasticity as a response to changes between particular environment sites and across all five sites, resulting in multiple plasticity measures for the same genotype. Despite a high overall correlation among these plasticity measurements, different QTLs were detected, indicating that these measures captured different aspects of plasticity with complementary information. Such differences in quantifying phenotype plasticity may be highly relevant in applications where the testing site and targeted site are clearly defined. In particular, when the mechanism of environmental factors interacting with the plastic QTL is known, an accurate prediction could be made on germplasm performances under various deployment environments at the GenBank level, facilitating precise breeding designs in the future.

Among the 23 traits, yield traits were more plastic than other traits and involved larger contributions from both temperature and day length, as well as a larger proportion of G-by-E interaction. A similar result has been reported in D’Andrea et al (2013)^46^. A possible explanation could be that yield traits were the results of combined effects from vegetative and reproductive growth with demonstrated contribution from both temperature and photoperiod^43,44^ that likely to be equally important, while flowering time was predominantly regulated by temperature^42^ with a relatively smaller contribution from photoperiod. Future studies are required to explore how differences in genetic architecture among traits cause such differences in phenotypic plasticity.

### The genetic architectures underlying trait mean and plasticity

In consistent with previous studies^25,47^, we found partial overlaps between QTLs associated with trait mean and plasticity. However, our interpretation is that when treating phenotype plasticity as a measure of change for one polygenic trait across environments, such overlap is expected. Besides this, we also expect that i) plasticity is polygenic as a result of the polygenic architecture for the trait itself at different environments, ii) the degree of overlap between QTLs underlying trait mean and plasticity may vary across studies due to detection power, and iii) QTLs with altered genetic effect among environments are more likely to impact the variability of plasticity. Taking DTT as an example, we detected 7 loci for DTT mean and 9 loci for DTT plasticity with four loci overlapping. The magnitude change (chromosome 5: 6,462,711 bp) or the sign change (chromosome 9: 35,126,793 bp) of QTL effects resulted in variability in DTT plasticity, providing support for the allelic sensitivity model^48^. Even though we did not detect the chromosome 9 QTL in DTT mean scan at a genome-wide significance, a moderate association was found at a lower significance threshold. In line with this, when aggregating the allelic effects of mean or plasticity QTLs not detectable at genome-wide significance, we found, as a group, they were significantly contributing to the variation of both mean and plasticity measurements (Figure S9, Table S10). Given the polygenic and dynamic genetic architecture of trait mean across environments reported here and in previous research^30,49^, there might be a tighter connection between the genetic regulation of trait mean and plastic than we have previously acknowledged.

### Plasticity QTL may have been subjected to directional selection during the breeding program

Previous studies showed that the highly selected region during maize adaptation to temperate climate explained less G-by-E variation than the selected region^12^ and the allele frequency of plasticity QTLs were changed between temperate and tropical lines^27^, suggesting that directional selection may have shaped their genetic diversity. Here, we explored whether the 93 plasticity QTLs were selected during intense artificial selection by evaluating their allele frequency changes using two collections of breeding materials, one collection from China that has predominantly been deployed in the 1960s, 1980s, and early 2000s, and a second collection from US before and after 2003^50^. We found that the allelic frequencies of 42 (45%) plasticity QTLs consistently changed from 1960s to 1980s and from 1980s to early 2000 in Chinese collection, and before and after 2003 in US collection (Figure S10). Such an agreement indicates that it is likely that these plastic QTLs were subjected to selection rather than genetic drift. However, many plastic QTLs, found here or in earlier researches^24,25^, were also contributing to the trait mean, further studies would be required to explore whether such changes are results of directional selection or simply consequences of selection on the trait mean that is correlated with the plasticity.

### Fine mapping the QTLs and the molecular basis of variation in phenotype plasticity

We found that a few QTL peaks, such as the QTL on chromosome 5: 6-7 Mb (Fig. 3A; Figure S4) and chromosome 8: 123-130 Mb (Fig. 3A; Figure S6) simultaneously associated with multiple traits means and plasticity measurements, possibly being a consequence of extended linkage disequilibrium (LD). Fine mapping the causal variants underlying each trait mean and plasticity QTL and distinguishing whether these signals were tagging one common signal simultaneously associated with multiple trait means and plasticity measures or they were multiple variants each associated with one measure but in tight LD with each other is a daunting task. Even though detailed analysis (Supplementary note) showed that a large proportion of the SNPs were tagging the same causal variants (Figure S4, S6), there seemed to be multiple independent association signals underlying the same QTL (Figure S4, S6) for a few mean or plasticity measures. For example, a detailed exploration on the chromosome 5 QTL showed that multiple statistical independent SNPs were tagging different combinations of multiple functional haplotypes (Figure S4), illustrating a complex genetic architecture involving allelic heterogeneity, multi-allelic, pleiotropy, and genotype by environment interaction at the same time. To pinpoint the causal genes in presence of such complexity, we applied gene-based test^51^ aggregating summary statistics on SNPs up/down stream of the annotated protein-coding genes, and detected 300 genes (Table S11), among which 24% were simultaneously associated with both mean and plasticity measurements (106 for mean, 122 for plasticity, 72 for both). Among these genes, the maize FT gene *ZCN8* was detected in both means and plasticity of flowering traits, while *ZCN18* was only associated with STI plasticity^52^. A benzoxazinone synthesis gene cluster including *bx1/2/3/8* on chromosome 4 was detected with association to the mean of ELW. Similar conditional effects also had been found in mutant and overexpression of multiple flowering genes in *Arabidopsis*, such as *PRR3* in circadian clock^53^, *PIF4* in ambient temperature pathway^54^, and *HXK1* in sugar pathway^55^. Although future experimental validations are required to validate the biological mechanism undying such variation, the validation of two candidate genes in our study suggests that the effect of genes on complex traits may in general be context-dependent.

## Conclusion

In summary, we showed that the genetic architectures of maize agronomic traits were dynamic across China’s major summer maize production zone with the genetic effects of many QTLs being either local or regional due to interaction with environmental factors, leading to changes in additive genetic variance, narrow sense heritability and variation in phenotype plasticity. The dynamic allelic effects of plasticity QTLs enable us to develop a GEAI complex trait prediction model with site-specific predictions and higher accuracy, opening a new possibility for future crop improvement. Our study thus provided novel insights into the dynamic genetic architectures of agronomic traits in response to changing climate and provided a GEAI model with site-specific prediction, paving a practical route to precision agriculture.

## Materials and methods

### Experimental design

We developed a Complete-diallel plus Unbalanced Breeding-derived Inter-Cross (CUBIC) population of 1404 maize inbred lines and surveyed their performance for 23 agronomic traits at five sites in China’s major maize production zone with longitudinal variation from E 114° 01’ at Henan (HN) to E 125° 18’ at Jilin (JL) and latitudinal variation from N 43° 42’ at HN to N 35° 27’ at HN. A detailed description of the development of this population was available in Liu et al^29^. Briefly, these inbred lines were derived from 24 elites representing 4 divergent heterotic groups with cycles of random mating, selection, and inbreeding^29^. In 2014, all inbred lines, each with five replicates, were planted at five sites, including Jilin Province (JL, N 43° 42’, E 125° 18’), Liaoning Province (LN; N 42° 03’, E 123° 33’), Beijing (BJ; N 40° 10’, E 116° 21’), Hebei Province (HB, N 38° 39’, E 115° 51’) and Henan Province (HN; N 35° 27’, E 114° 01’). Twenty-three agronomic traits, including 6 phonology traits, 8 plant architecture traits, and 9 yield traits were phenotypically evaluated. Except for six flowering traits that were scored as the median values of replicated lines, all the remaining traits were scored as the means among replicates (Table S12). Three environmental variables, including daily highest temperature (TemH), daily temperature difference (TemD), and day length (DayL), were extracted from local weather stations (http://data.sheshiyuanyi.com/WeatherData/). All the 1404 lines were re-sequenced and called genotypes were available for download from Liu et al^29^. Totally 6.6 M SNPs with MAF > 0.03 and LD =< 0.9 in 100 kb sliding window were retained for downstream analysis.

### Partition the phenotypic variance into contributions from genotype, environment factors, and their interactions

The phenotypic variance was partitioned into contributions from genotype (G), genotype-by-environment (G-by-E), and residual (environment; E) by fitting the following model:

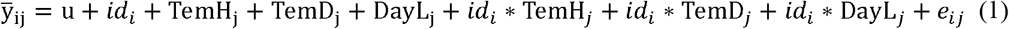

This model was fitted for each of the 23 traits one at a time. ȳ_ij_ is the trait mean/median of individual i (i= 1…n, n = 1404 is the number of individuals) at site j (j = 1…q; q=5, corresponds to the number of sites); idi is the line id (genotype) coded as factor; TemH_j_, TemD_*j*_, and DayL_j_ are three environmental variables representing the daily highest temperature, daily temperature difference, and day length at site j, respectively. These environmental factors were coded as numeric, assuming a linear relationship with the phenotypic measurements. *id_i_*\* TemH_j_, *id_i_*\* TemD_j_, and *id_i_*\* DayL_j_ are the interaction terms (G-by-E) between a particular line (genotype) and the corresponding environmental factors (environment). The relative contributions to the total phenotypic variance from G and G-by-E were estimated by their respective sum of squares (Sum of Square for id is calculated as 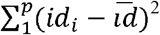 and Sum of Square for the interaction terms id* *E* are calculated as 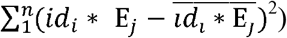, where E stands for TemH, TemD, and DayL.

### Estimating the narrow-sense heritability, additive variance, and genetic correlations

A linear model was used to estimate the narrow-sense heritability for all the 23 traits measured at each of the five sites.

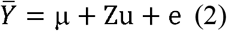

Here, *ȳ* is a vector of trait mean/median of each individual (genotype) at each tested site. *e* is the normally distributed residual. μ is a column vector of 1’s to represent the population mean, and u is a random effect vector of the breeding values for the 1404 individuals. Z is the corresponding design matrix obtained from a Cholesky decomposition of the kinship matrix G, estimated using the genome-wide markers using GCTA^56^. The Z matrix satisfies ZZ’=G, therefore, that is normally distributed 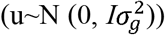. e is the residual variance with 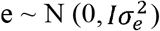. The narrow-sense heritability of fitted phenotype was calculated as the intraclass correlation 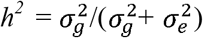. AI-REML implemented in GCTA was used to obtain these estimates^56^. The additive genetic variance was then estimated as the variance of Y (Var_Y_) times *h^2^*.

Similarly, a bivariant mixed model was fitted to obtain estimates of the genetic correlation between measurements obtained from two individual sites. Ten models were thus fitted to obtain all the pairwise genetic correlations among five sites. *ȳ*, μ and u from the model (2) were updated to an n×2 matrix, with n being the number of individuals and each column vector representing measurement obtained from a particular site. This model was fitted using the *reml-bivar* module^57^ implemented in the GCTA software^56^ and details of this model were available in Lee et al^57^.

### Quantification of the phenotypic plasticity for the 23 agronomic traits

Since all the 1404 maize lines were phenotyped for 23 agronomic traits across five sites, we quantified and studied the genetics of maize complex trait plasticity in response to longitudinal and latitudinal environmental variation. Here, the phenotypic plasticity was classified into two categories (Figure S2B-E). The first category is overall plasticity describing plasticity across all the studied environments, while the second category-specific plasticity is more unique to certain pairs of sites, which only captures the plasticity across two environments. The motivation underlying such classification is that some individuals are more robust across most of the studied sites except only one or a few sites, while other individuals are plastic among most of the sites.

One metric, pairwise difference in phenotypic value between two sites, was used to quantify the specific plasticity (Figure S2A). Using DTT measured at JL and HN as an example, the differences in measured DTT values for all individuals 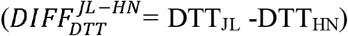 would describe the specific plasticity between HN and BJ (Figure S2B)^31^. In addition, four additional approaches were used to quantify the overall plasticity (Figure S2C-E). First, the principal component analysis (PCA) was used to quantify the overall plasticity. The influence of the phenotype measures at individual sites on the principal components (PCs) can be captured in the loadings^58^ (Figure S2C). As the second PC (PC2) captures more variation in overall plasticity, we consider PC2 as a measure for overall phenotypic plasticity. Second, the across environment variance (VAR) of the rank transformed phenotype proposed in Vanous et al., 2019 was used (Figure S2D), and the coefficient of variance (CV) ^7^ was also used to account for the mean difference. The fourth score for the overall plasticity (FWR) applies Finlay–Wilkinson Regression^32,59^ to partition the phenotype into two components, one is constant across environments and another responds dynamically to environmental changes. Using the linear mixed model, the phenotype of each line is partitioned into these two components and the plasticity component is used as a measurement of plasticity. In total, the described approaches resulted in 14 measurements of phenotypic plasticity (abbreviated as DIFF, PCA, VarR, CV, and FWR). Altogether, these three metrics yield 14 plasticity measurements for each trait.

### Genome-wide association analysis for the trait mean/median and plasticity measurements

To detect genetic polymorphisms underlying variation of agronomic trait mean and plasticity, we fitted the following linear mixed model:

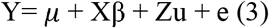

where Y, *μ*, Z, u, and e are the same as has been defined in the model (1). X is a matrix containing the genotype of the tested SNP (coded as 0/2 for minor/major-allele homozygous genotypes, respectively). β is a vector including the estimated additive allele-substitution effect for the tested SNP. First, a genome-wide analysis (GWA) across all genotyped SNPs was conducted using GEMMA^60^. A subsequent conditional analysis was performed where all the top associated SNPs (the SNPs with the highest P value from each association QTL from the initial GWA scan) were included as covariates in the design matrix X to screen for additional association signals. This conditional analysis was repeated until no more SNPs were above the significance threshold. This conditional analysis was implemented in *cojo* module from GCTA^61^. The linkage disequilibrium (LD) was high in this population, making Bonferroni correction assuming all tested markers were statistically independent too conservative. Therefore, we estimated the effective number of independent markers (Me)^62^ and derived a less conservative genome-wide significance threshold following 0.05/Me (1.53 x 10^-8^ equivalent to −Log_10_^p^ = 7.81).

### Colonization test separates linkage from pleiotropy at regions where multiple signals were associated with multiple traits

At the same genomic region, multiple association signals, each associated with one or multiple traits, were colocalized. Since the level of LD between the lead SNPs is very low, we could not directly tell whether multiple independent signals, detected in multiple scans and physically close to each other, are from one association signal simultaneously associated with multiple scans (pleiotropy), or multiple associations each associated with one scan but in tight LD with each other. To distinguish this, we performed a multi-trait colocalization analysis (Supplementary note). This method estimates a posterior probability of whether multiple traits are sharing a common causal variant using summaries statistics from each trait^63,64^. We first binned the genome into 1 Mb bins. Scans with independent SNPs that fall into consecutive bins were aggregated and tested for colocalization using the *hyprcoloc* R package^63,64^. Given the complex population history (multi-parental) and a limited number of recombinations, some of the statistically independent SNPs were very close to each other. To make a comparison among the 4 types of measurements, we arbitrarily grouped SNPs less than 1Mb to a single QTL.

### Gene-based test to prioritise candidate genes

The LD was too extensive to directly pinpoint the genes underlying the associated loci. We, therefore, applied a set-based analysis that aggregates summary statistics from all the variants 50 kb up/downstream of the tested gene to obtain one p value to represent the significance of a particular gene. The summary association statistics, including effect sizes, standard errors, minor allele frequencies, and sample size, were first extracted from the GEMMA association output, and subsequently inputted to *fastBAT* module in GCTA^65^. And 39,155 genes, annotated in the B73 reference genome version 3 were used to bin the summary statistics to perform the set analysis^51^.

### Testing for genotype by environment interaction of detected QTLs

We tested the interaction between QTLs associated with each of the 23 traits in at least one of the five sites, one QTL and one trait at a time. This was done by fitting the model below:

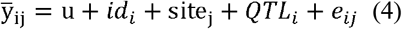

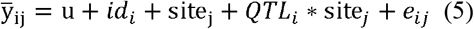

This model was fitted for each of the 23 traits one at a time. ȳ_jj_ is the trait mean/median of individual i (i= 1…n, n = 1404, number of individuals) at site j (j = 1…q; q=5, corresponds to the number of sites); id_i_ is the line id (genotype) coded as a factor; site_j_ is a vector of characters representing the site where the measurements were made. *QTL_i_* is the genotype of id_i_ at the testing QTL, and *QTL_i_* * site_j_, is the interaction terms (G-by-E) between a particular QTL and the sites (environments). A likelihood ratio test comparing the model with (Model 4) and without (Model 5) the interaction between sites was performed to calculate p values. The significance threshold was derived as 0.05 dividing the number of tests (0.05/143= 3.49×10^-04^)

### Experimental validation of maize flowering genes

Knock-out lines of *ZmTPS14.1* and *ZmSPL6* were generated using a high-throughput genome-editing system^35^. In brief, line-specific sgRNAs were filtered based on assembled pseudo-genome of the receptor KN5585. The Double sgRNAs pool (DSP) approach was used to construct vectors. The vectors were transformed into the receptor KN5585. The genotype of gene-editing lines was identified by PCR amplification and Sanger sequencing using target-specific primers (Table S13). The phenotype of knock-out lines and wild type were investigated in Jilin (Gongzhuling, Jilin province, N 43° 30’, E 124° 49’) and Hainan (Sanya, Hainan Province, N 18° 34’, E 108° 43’).

### Time-course transcriptome

B73 seeds were planted at two conditions, long-day condition (14 hours light and 10 hours dark) and short-day condition (8 hours light and 16 hours dark). Leaf tissues were harvested at 3 time points in one day at stage V4 (Vegetative 4, four fully extended leaves). Eighteen samples (2 conditions × 3 time points × 3 replicates) were RNA-sequenced by Hiseq3000. Low-quality reads were filtered out by trimmomatic^66^. STAR^67^ was used to align the RNA-seq reads to the reference genome. HTSeq^68^ was used to obtain gene-level counts from the resulting BAM files. Genes with significant expression changes were detected by ImpulseDE2^69^.

### Estimating the contribution from mean and plasticity QTLs to the variation of mean and plasticity measurements

We quantified the contribution from mean and plasticity QTLs to the variation of trait mean and plasticity by fitting the following models.

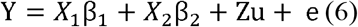

Here, Y is a vector of length n (n =1404), representing the trait mean or plasticity measurement. The joint contributions from mean and plasticity QTLs were modelled in and *X_2_β_2_* where X1 and X2 are the design matrixes β_1_ and β_2_ are the corresponding effect sizes. Z, u and e is the same as defined in model 3. Contributions from mean and plasticity QTL were then calculated with 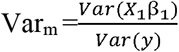 and 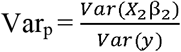.

### Forecasting the site-specific performance of the 23 traits

We fitted the following model to predict the performance of each site for the 23 traits one at a time.

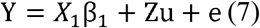

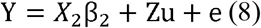

Here, Y is a vector of length n*p (n =1404, the number of individuals; p = 5, the number of sites; n*p = 7020), representing the trait means measured at five sites. u is a vector of length n*p, representing the breeding value of the n maize line, and e is the randomly distributed residual with length n*p. The Z matrix satisfies ZZ’=G ⊗ *I*, where G is the identity by state (IBS) matrix and I is a diagonal matrix of pxp. *X_1_* is a design matrix with one column of 1 representing column mean and additional 4 columns representing the environmental effects from the remaining 4 sites, and β_1_ is a vector of corresponding effect sizes. *X_2_* includes all the columns from *X_1_* and additional columns with genotypes of the k QTL associated with the mean and plasticity measures of the tested trait, and additional columns representing the interaction between the k QTL and the five sites, capturing the effects from QTL by environment factor interaction. The fitted values from model 7 were referred to as GBLUP predictions while the fitted values from model 8 were referred to as GEAI predictions. These models were fitted using rrBLUP^70^ package in R (https://www.R-project.org/). In the first case, we evaluated the predictability on untested lines at any of the five sites by using all the lines phenotyped across the five sites using 5-fold cross-validation. Each time, 80% of the lines were randomly sampled and used to predict the remaining 20% lines. In the second case, we simulated a serial of more challenging breeding designs, in which only a core set of lines (10% - 70%) were phenotyped across five sites and the interest was to predict the performance of unphenotyped lines at each site. Each time, a core set of lines were randomly sampled and the remaining lines were divided into 4 sets and were randomly assigned to one of the remaining 4 sites, whose phenotypes were masked as NA and unassigned environments. Accuracies were estimated as the regression r^2^ between measured and predicted phenotypes.

## Supporting information

CUBIC_GE_S_Figs

CUBIC_GE_S_Note

CUBIC_GE_S_Tabs

## Supplementary materials

Figure S1-S10, Table S1-S13, and note were available in supplementary files

## Author Contributions

Conceptualization, J.Y., M.J., Y.Z., H.L.; methodology, Y.Z., M.J., H.L.; formal analysis, Y.Z., M.J.; investigation, Y.Z., M.J., H.L.; data collection and curation, X.L., J.G., Y.Y., Z.L., J.Z., X.W., F.Q., M.J., Y.Z., H.L.; writing original draft preparation, Y.Z., M.J.; writing reviewing and editing, M.J., Y.Z., H.L., T.G., Y.X., J.Y.; visualization, Y.Z., M.J, Y.J; supervision, J.Y.; funding acquisition, J.Y., Y.Z., H.L., X.L. All authors have read and agreed to the published current version of the manuscript.

**The authors have no conflicts of interest to declare.**

## Acknowledgments

We would like to thank the Swedish National Infrastructure for Computing (SNIC) for their support in computation resources through High Performance Computing Centre North (HPC2N, SNIC 2020/9-117) and the Uppsala Multidisciplinary Centre for Advanced Computational Science (UPPMAX).

