## Supplementary material for "Complex genetic architecture underlying the plasticity of maize agronomic traits": CUBIC_GE_S_Figs

Supplementary figures for **‘Complex genetic architecture underlying the plasticity of maize agronomic traits’**

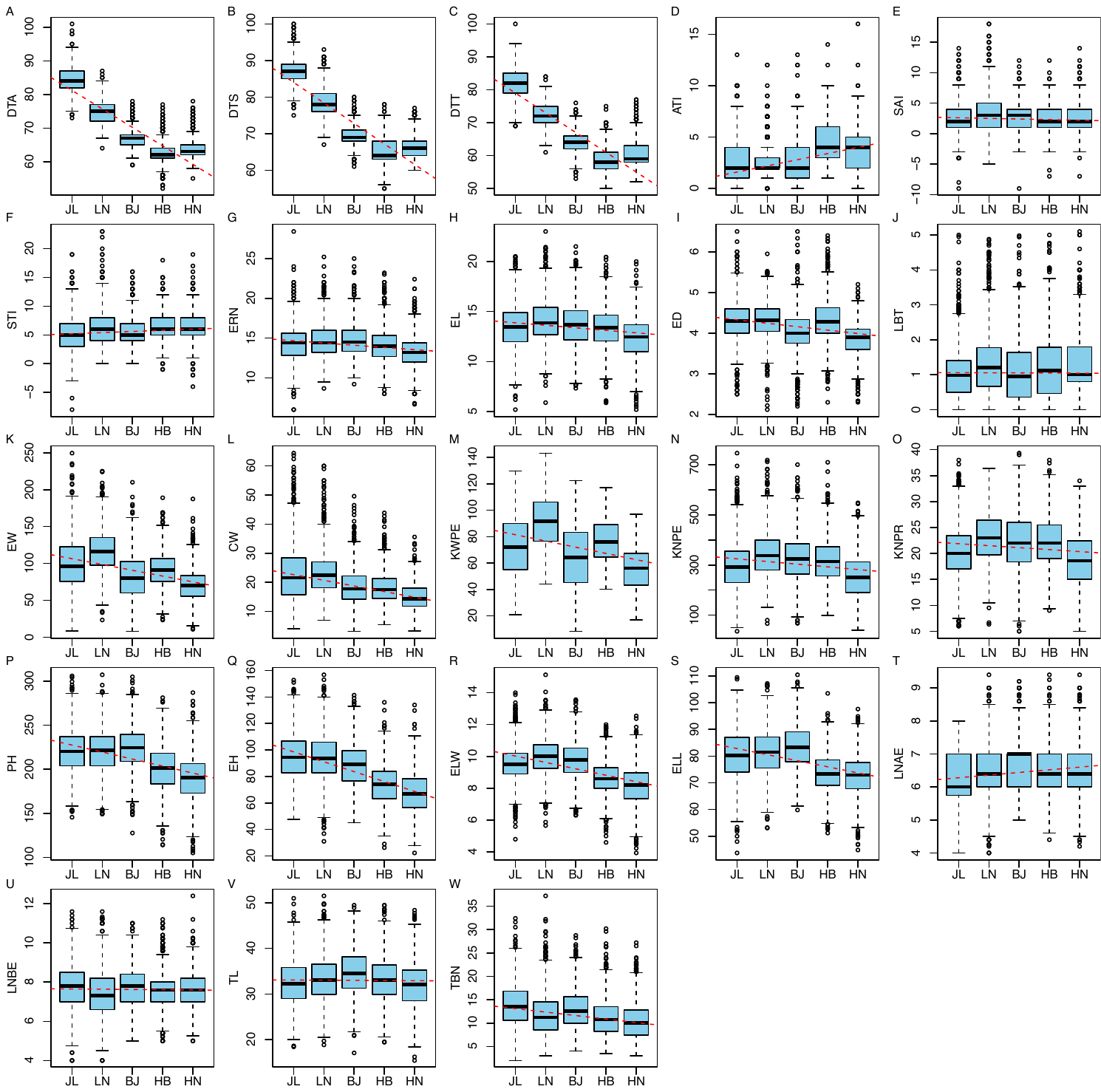

**Figure S1.** Boxplot of the phenotypic measurements for 23 maize complex traits at the five different sites.

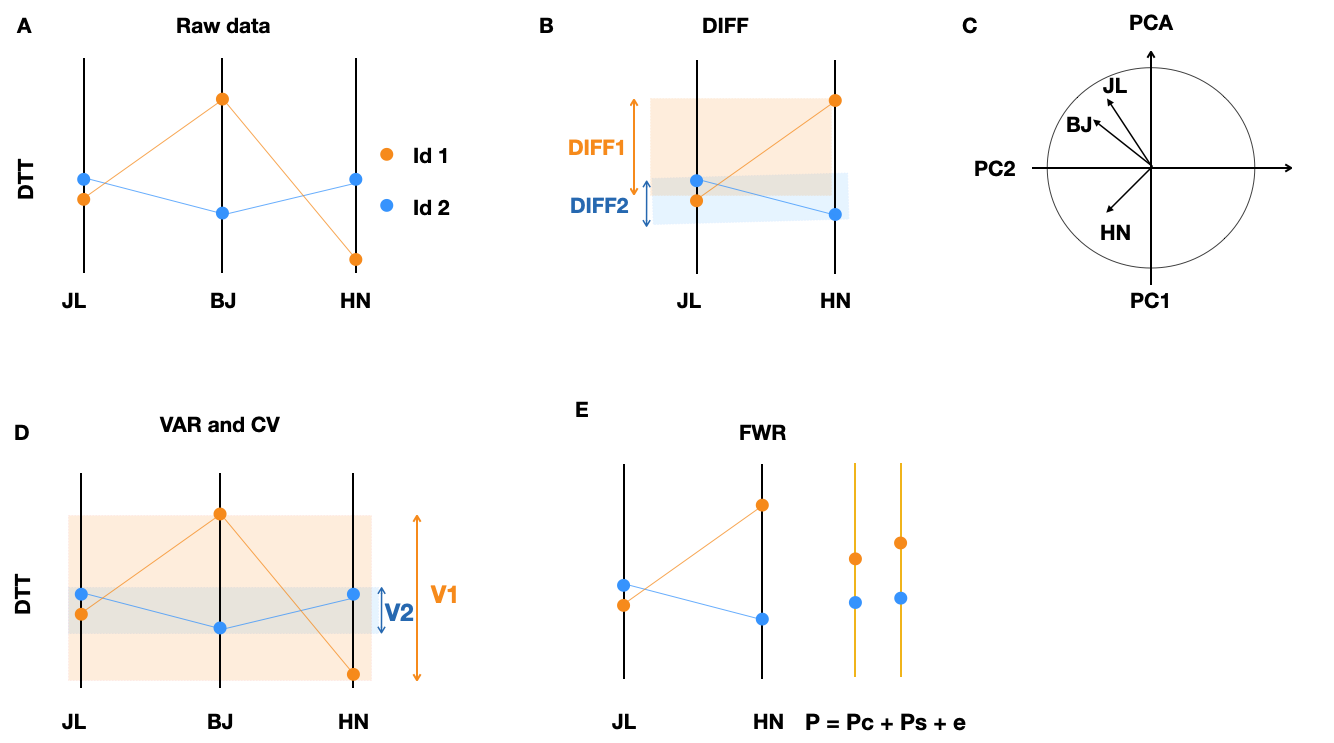

**Figure S2.** Schematic illustrations of the approaches to quantify the phenotype plasticity of DTT. **A).** A conceptual illustration of phenotypic measurements across 3 sites and phenotype plasticity. Each bar represents an environment and the average phenotype of two maize line (genotypes, id1 and id2) are illustrated using orange/blue dots. Average DTT for the two lines (horizontal blue and orange lines) varied across JL, BJ and HN (vertical black lines). The DTT for one line (orange) varied more (more plastic) in response to the environmental changes than the other line (blue). **B).** Pairwise difference in DTT between JL and HN quantifying the specific plasticity between these two sites. All the 10 possible combinations among the five sites were used in our study, but only JL and HN was selected here for simplicity. Four measures of overall phenotype plasticity across all environments (JL, LN, BJ, HB, HN) were used and schematically illustrated in (C–E). **C).** A variable loading plot showing the contributions by the three sites (only JL, LN, HN were visualised for simplicity) to the first two principal components (PC1 and PC2). PC2 explaining the most variance was used as measures of plasticity. The second overall measure (VarR, **(D)**) quantifies plasticity as the across environment variance of rank transformed DTT. The third overall measurement (FWR, **(D)**) partitions DTT into two components, one (Pc) that is constant across environments and another (Ps) that varies across environments. Here, DTT measured from two environments, JL and HN were illustrated on the left, for simplicity.

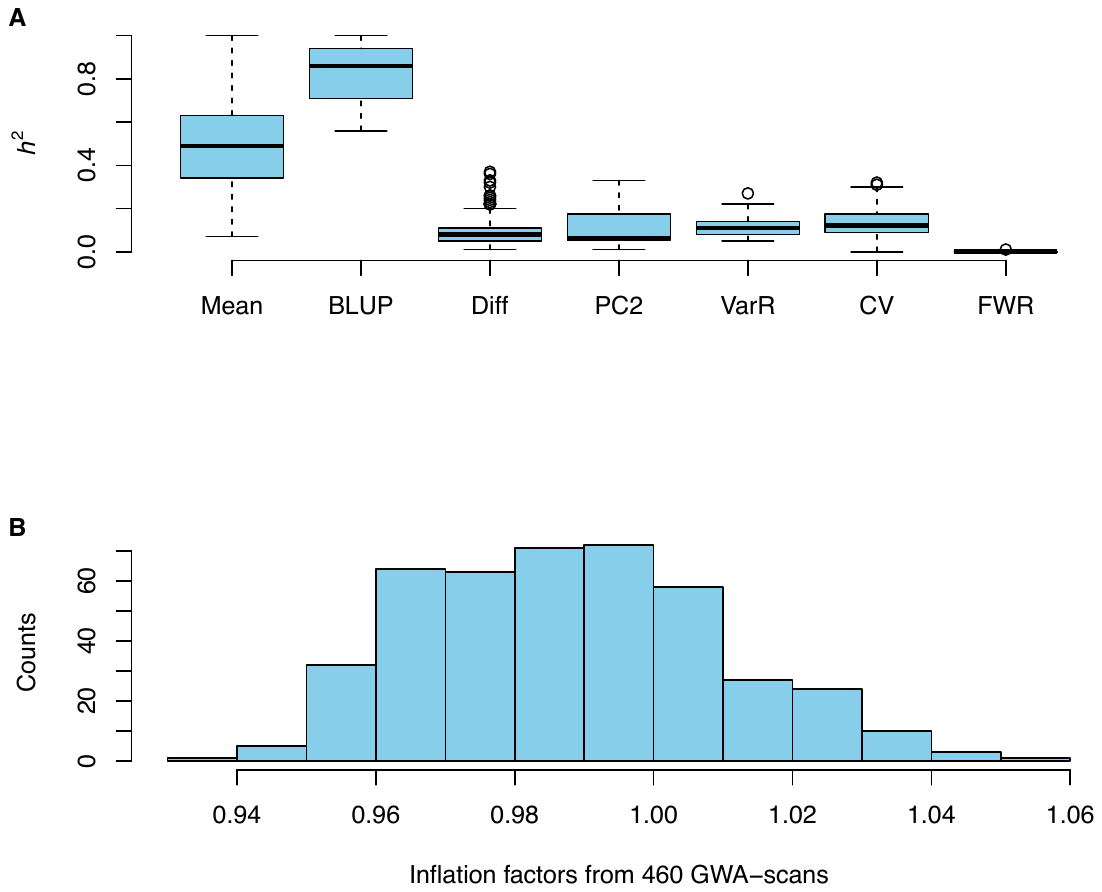

**Figure S3.** Illustration of the narrow sense heritability **(A)** and inflation factors **(B)** of the 460 GWA-scans.

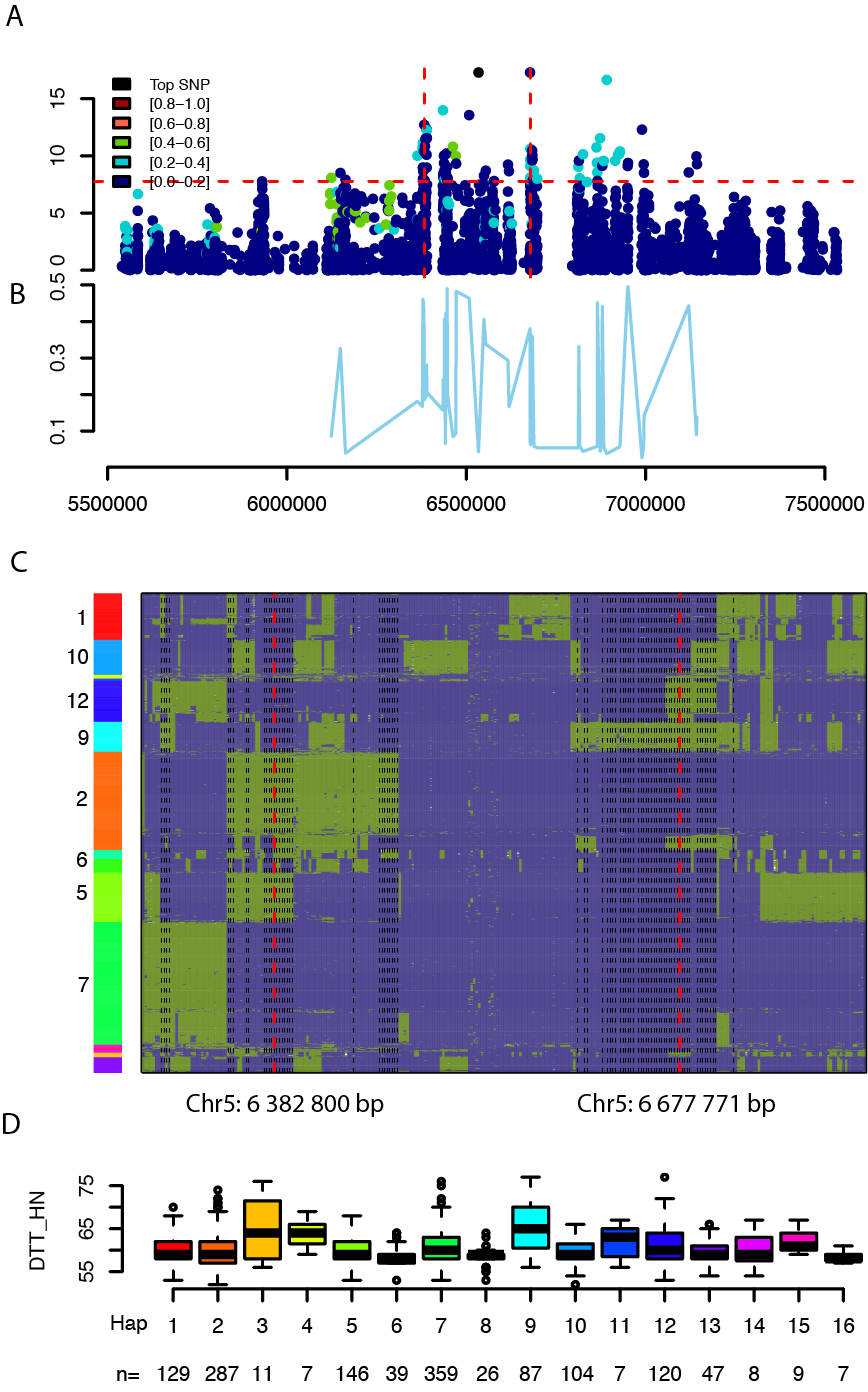

**Figure S4.** A detailed exploration of the association signal on chromosome 5. **A).** Manhattan plots for SNPs, from 5.5 Mb to7.5 Mb on chromosome 5, associated with DTT_BJ_. **B).** Allele frequencies for SNPs with p value below the genome wide significant threshold. **C).** Haplotype structure at this region. Each cell represents the genotype of one individual (row) at a particular SNP (column), with deep blue and olive green represents two alternative genotypes. The index to the left highlights the haplotypes defined by clustering the genotype matrix. Significant SNPs were highlighted using dashed lines, where the two independent SNPs (chromosome 5:6 462 711 bp and 6 382 800 bp) detected in conditional scan were highlighted red and others were in black. **D).** Genotype to phenotype map with the 16 haplotypes defined from hierarchical clustering of SNPs from this region and DTT_HN_.

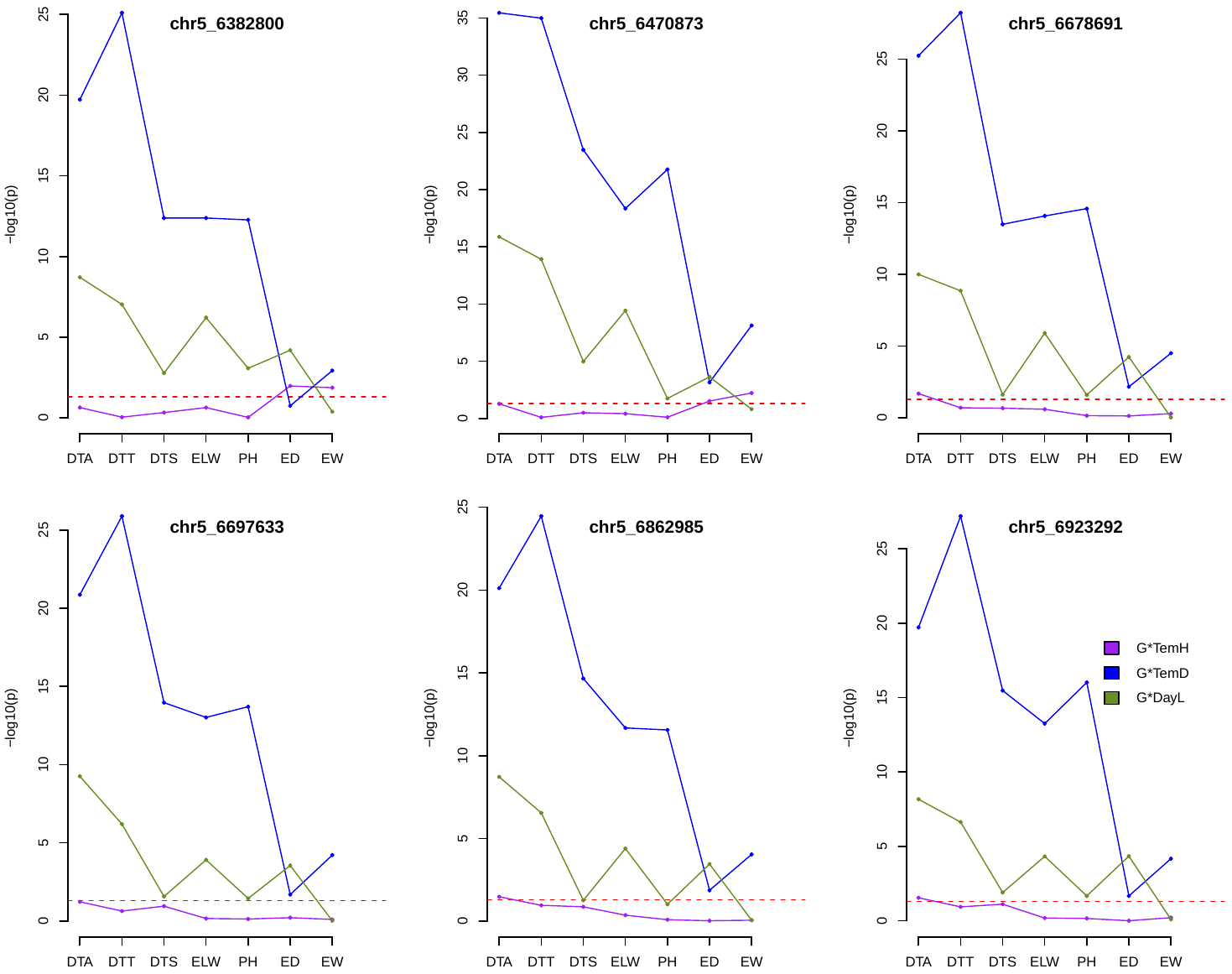

**Figure S5.** Illustration of the p-values testing the interaction between the 6 SNPs from cluster 3 and various environmental factors.

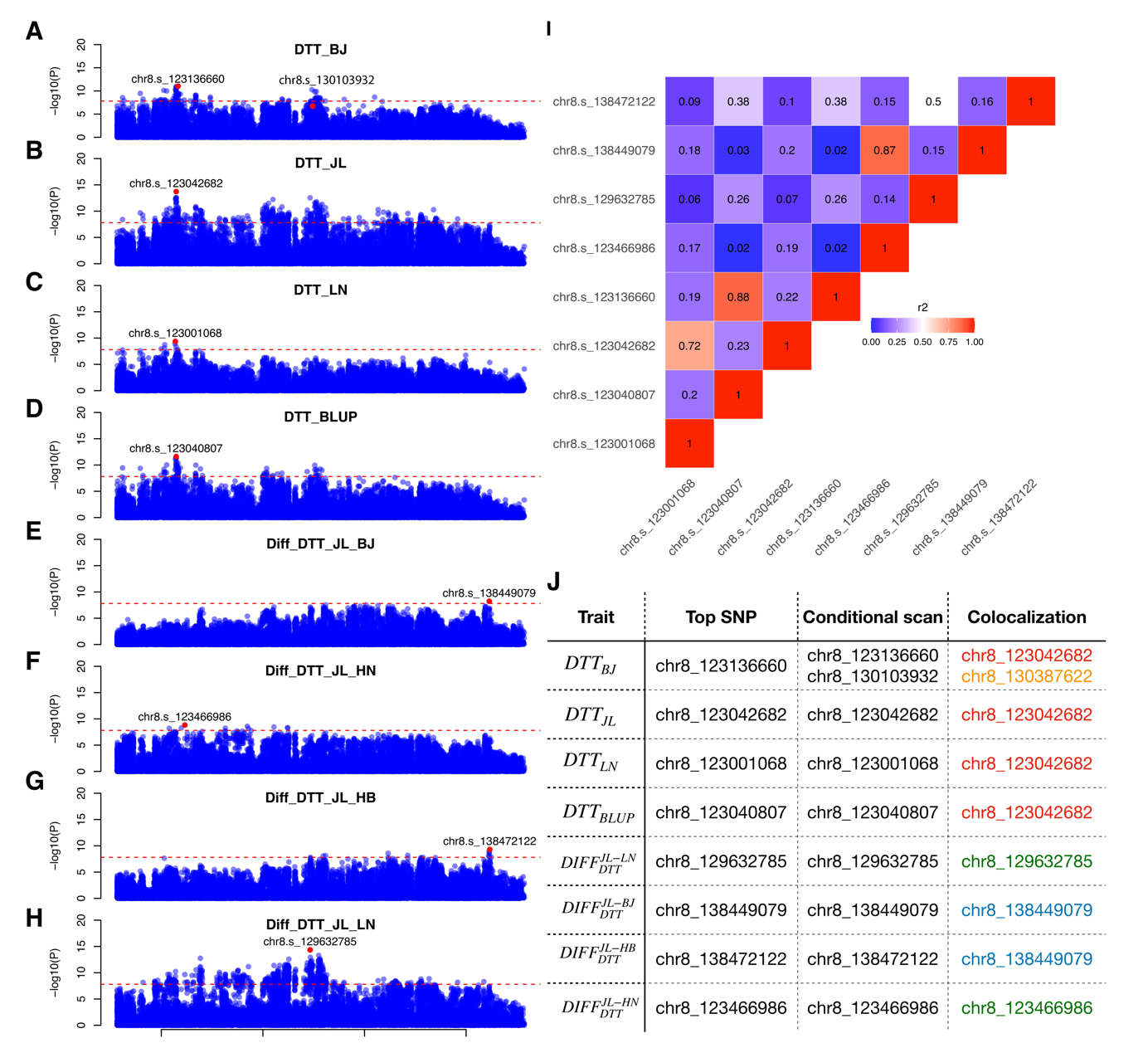

**Figure S6. A-H)** Manhattan plot for the association signal from 120 Mb to 140 Mb on chromosome 8 for the 8 scans with significant associations. The name of the scan is labelled in the middle of the panel. **I).** LD heatmap of the r2 value of the top associated SNPs. **J).** The associated SNPs from standard GWA scan, conditional scan and colocalization scan. Shared loci are marked in the same colour, except for green colour indicating colocalization analysis did not bring any information.

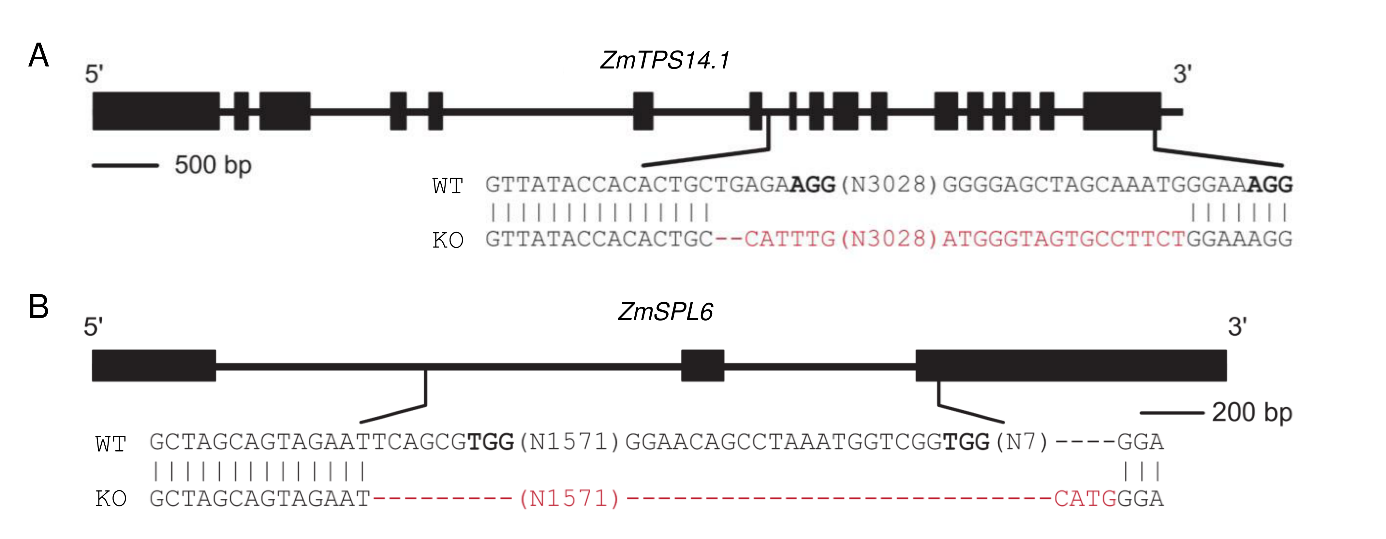

**Figure S7.** Gene structure and targeted sequences of *ZmTPS14.1* (**A**) and *ZmSPL6* (**B**) in wild type and knock-out lines. Two sgRNAs were designed and a large inversion and deletion were caused in *ZmTPS14.1* and *ZmSPL6*, respectively.

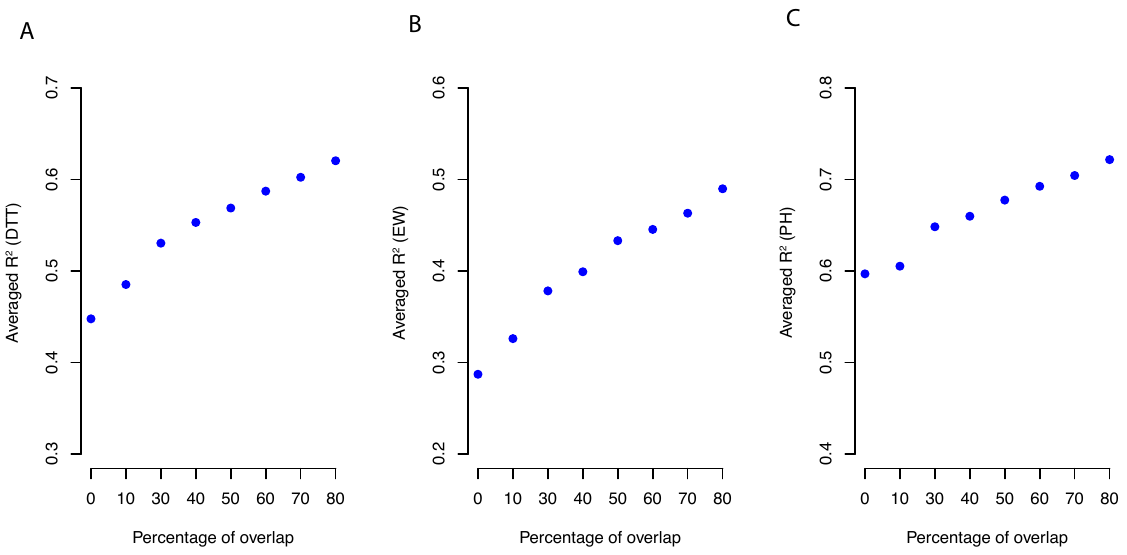

**Figure S8.** Averaged prediction accuracy across 5 sites using various degree of phenotyped overlapping lines from 5 sites for **A)** DTT, **B)** EW and **C)** PH.

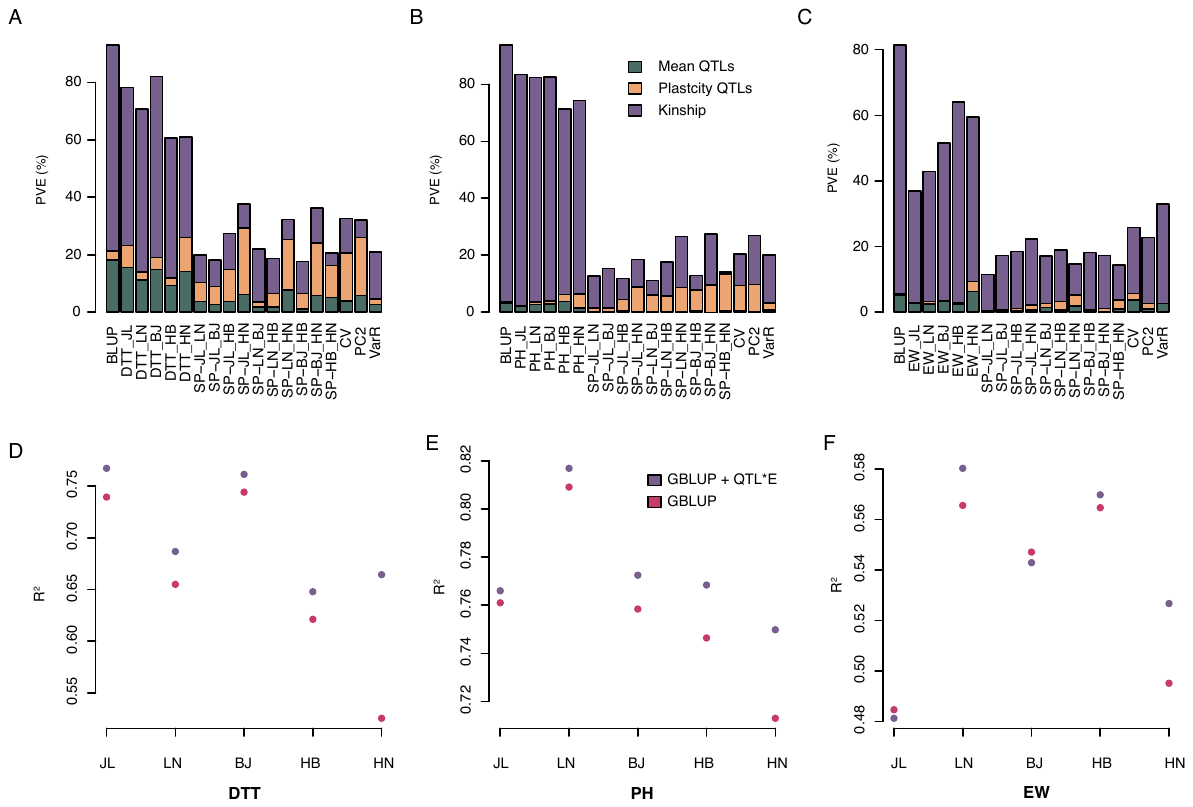

**Figure S9.** Contribution from mean QTLs, plasticity QTLs and kinship to the variation of the mean and plasticity measurements for **A)** DTT, **B)** PH and **C)** EW. Each vertical bar represents a trait mean or plasticity measurements with the corresponding trait name labelled in x-axis. The coloured segments within each bar represent the contribution from mean QTLs, plasticity QTLs and kinship as indicated in the legend. The height of the bar is proportional to the variance explained (PVE) by the corresponding variance component.

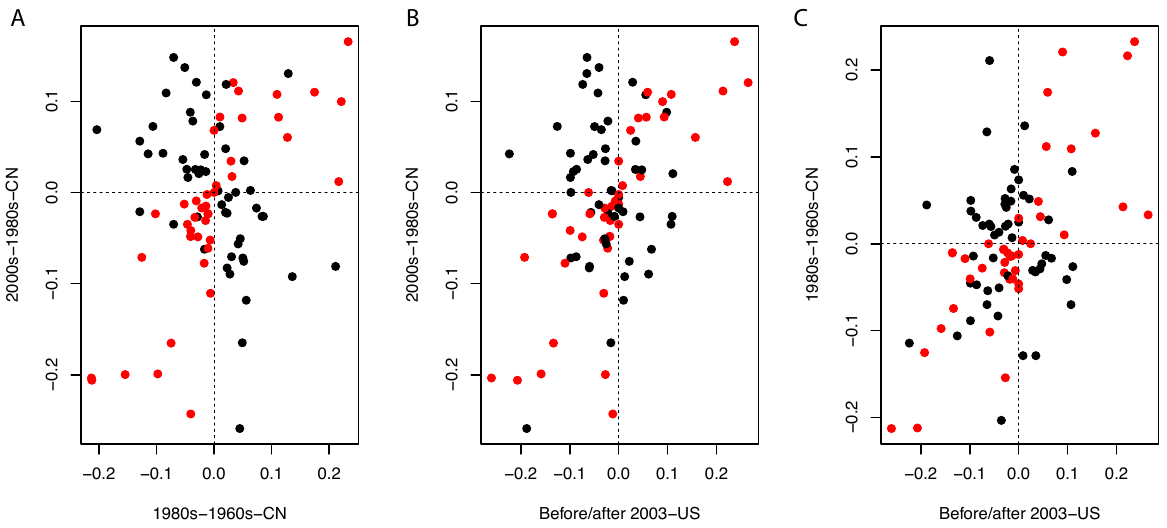

**Figure S10. Allelic frequency change of the 93 plasticity QTLs in a collection of breeding materials from China that has predominantly been deployed in the 1960s, 1980s and early 2000s, and a second collection of breeding materials from US line before and after 2003.**

**A).** Scatter plot of allele frequency difference between 1980s-1960s-CN lines (x-axis) and 2000s-1980s-CN lines (y-axis). Each dot represents an allele, the value on x-axis is the allele frequency difference from Chinese lines deployed in 1980s and1960s, and the value on y-axis is the allele frequency difference from Chinese lines deployed in 2000s and 1980s. **B).** Scatter plot of allele frequency difference between US lines Before/after 2003 (x-axis) and 2000s-1980s-CN lines (y-axis). **C).** Scatter plot of allele frequency difference between US lines Before/after 2003 (x-axis) and Chinese lines deployed in 1980s and1960s (y-axis).
