## Supplementary material for "Complex genetic architecture underlying the plasticity of maize agronomic traits": CUBIC_GE_S_Note

Supplementary note for ‘**Complex genetic architecture underlying the plasticity of maize agronomic traits**’

***Complex genetic architecture involving allelic heterogeneity, pleiotropy and genotype by environment interaction for the locus on chromosome 8***

As the linkage disequilibrium was very extensive to directly pinpoint the causal variant/genes, we attempted to selected the independent association signals by applying conditional analysis and colocalization analysis. Taking DTT for an example, for the peak on chromosome 8, 2 independent association signals, one on 123,136,660 bp and a second one on 130,103,932 bp, were detected for DTT_BJ_ using conditional scan (Figure S6A), but only one of them (Figure S6B-H ,chromosome 8: 123,001,068 bp for DTT_LN_, chromosome 8: 123,040,807 bp for DTT_BLUP_, chromosome 8: 123,042,682 bp for DTT_JL_) was detected for the remaining DTT-scans, suggesting the presence of multiple independent associations with variable genetic effects under different environments. The level of linkage disequilibrium (LD) between the independent SNPs, detected in separate DTT-scans, was very low (median r^2^ = 0.18; Figure S6I), making it very challenging to distinguish whether these signals were tagging one common signal simultaneously associated with multiple DTT-scans, or they were multiple variants each associated with one scan but in tight LD with each other. To separate these two scenarios, we performed colocalization analysis (see details in Materials and Methods). Besides a few cases, the majority of the top associated SNPs were likely tagging one common signal simultaneously associated with multiple scans. Taking the locus on chromosome 8 as an example (Figure S6J), the three variants, chr8: 123,136,660 bp associated with the variation of DTT_BJ_, chr8: 123,001,068 bp associated with the variation of DTT_LN_ and chr8: 123,040,807 bp detected for DTT_BLUP_ scan, were colocalised with the SNP (chromosome 8: 123,042,682 bp) associated with DTT_JL_ (Figure S6J). There are three additional independent signals, being either unique to one scan or shared by two DTT-scans (Figure S6J), suggesting the presence of multiple independent signals at this genomic region with variable effects on different DTT measurements.

***Multiple haplotypes underlying the association peak on chromosome 5***

We detected 7 statistically independent SNPs underlying the association peak on chromosome 5 associated with DTT_HN_. As these 7 SNPs were only 540 kb apart from each other, we took a detailed exploration. Figure S4A illustrated the Manhattan plot of SNPs associated with DTT_BJ_ from 5.5 Mb to 7.5 Mb on chromosome 5, while Figure S4B showed the minor allele frequency (MAF) for all the SNPs above genome wide significant threshold at this region. Two SNPs (chromosome 5: 6,462,711 bp and chromosome 5: 6,382,800 bp) were detected in the conditional scan, suggesting the presence of two statistically independent signals (Marked using dashed vertical red lines in Figure S4A). Notably, the MAF of these SNPs varied from 0.03 to 0.5, indicating that they were likely tagging different causal variants. This hypothesis was supported by the observation of 16 haplotypes, with statistically significant genetic effects (Figure S4D; P= 2.78 x 10^-4^ from ANOVA), obtained by clustering the genotypes of all the 1404 individuals within this region (Figure S4C), The significant SNPs detected from the single marker GWA scan were indeed contrasting different groups of haplotypes causing SNPs with distinctive allele frequencies being statistically significant.
